# A frontosensory circuit for visual context processing is synchronous in the theta/alpha band

**DOI:** 10.1101/2023.02.25.530044

**Authors:** Georgia Bastos, Jacob T. Holmes, Jordan M. Ross, Anna M. Rader, Connor G. Gallimore, Darcy S. Peterka, Jordan P. Hamm

**Author notes:** Corresponding author: Jordan Hamm, 813 Petit Science Center, 100 Piedmont Ave, Atlanta, GA 30303.

## Abstract

Visual processing is strongly influenced by context. Stimuli that deviate from contextual regularities elicit augmented responses in primary visual cortex (V1). These heightened responses, known as “deviance detection,” require both inhibition local to V1 and top-down modulation from higher areas of cortex. Here we investigated the spatiotemporal mechanisms by which these circuit elements interact to support deviance detection. Local field potential recordings in mice in anterior cingulate area (ACa) and V1 during a visual oddball paradigm showed that interregional synchrony peaks in the theta/alpha band (6-12 Hz). Two-photon imaging in V1 revealed that mainly pyramidal neurons exhibited deviance detection, while vasointestinal peptide-positive interneurons (VIPs) increased activity and somatostatin-positive interneurons (SSTs) decreased activity (adapted) to redundant stimuli (prior to deviants). Optogenetic drive of ACa-V1 inputs at 6-12 Hz activated V1-VIPs but inhibited V1-SSTs, mirroring the dynamics present during the oddball paradigm. Chemogenetic inhibition of VIP interneurons disrupted ACa-V1 synchrony and deviance detection responses in V1. These results outline spatiotemporal and interneuron-specific mechanisms of top-down modulation that support visual context processing.

## Introduction

The brain uses context when processing sensory information to support perception and guide behavior. Context can include perceived patterns about the environment – both spatial and temporal information – along with any assumed regularities about which stimuli may be predictable versus unexpected. In early cortical regions, contextual modulation of sensory evoked responses serves to sharpen perception and streamlines information processing to guide learning and behavior. Understanding how these neuronal circuits incorporate and process contextual information is therefore paramount.

One popular generalized framework for understanding and studying context processing in mammalian neocortex is “predictive processing”^1–3^. Herein, the inferred causes of sensory inputs – or predictive models of the environment – are encoded in cortical areas hierarchically upstream from primary sensory cortices, such as secondary/tertiary visual, parietal, or pre-frontal cortices^2, 4^. Such higher areas of cortex send inputs to lower areas representing that include “predictions” about future sensory input. These inputs can effectively modulate sensory processing, suppressing cortical responses to stimuli which are expected by the predictive model encoded in the higher brain area. Stimuli which deviate from this model elicit “prediction errors” in sensory cortex – typically large neuronal responses to the stimuli in a subset of neurons – which then propagate to higher areas to update the predictive model of the environment (and beliefs about the underlying causes of sensory input)^5–7^.

Evidence in support of this predictive coding model of sensory processing in the cortex spans various sensory modalities in multiple mammalian species, from humans to non-human primates^8, 9^, cats, and rodents^10^. Furthermore, recent studies have employed mice to study the cell and circuit mechanisms of predictive processing^8,11–14^, often using primary visual cortex (V1) during a visual “oddball” paradigm as a model^11,14^. Oddball paradigms are simple and widely used sensory stimulation paradigms for studying basic predictive processing. An oddball sequence involves a repeated stimulus (the “redundant”) presented rapidly (≈1.1 Hz) and interspersed by a rare “deviant” stimulus (the “oddball”). In V1, evoked neuronal responses to the redundant stimulus are attenuated, while responses to the deviant stimuli are augmented^11,14,15^. This augmented activity, termed “deviance detection (DD)”, may represent a basic form of cortical “prediction error,” indexing a deviation of current sensory data from the expected input. Our recent study showed that only a subset of pyramidal neurons (PYRs) in V1 exhibit DD and that these neurons are enriched in layer 2/3 of cortex^14^ (L2/3), consistent with theoretical models of “prediction error” signal generation in cortical microcircuits^7^ and empirical reports of visuomotor prediction errors^12^.

Additionally, this work showed that top-down input to V1 from a prefrontal region, the anterior cingulate area (ACa), is necessary for the production of DD responses (i.e., prediction errors) during the oddball paradigm^14^. Interestingly, the axon terminals of ACa neurons projecting to V1 did not themselves exhibit DD responses, but rather were active during all phases of the oddball paradigm. This suggests that DD responses present in V1 are not simply inherited from top-down ACa drive but, rather, that they arise indirectly from ACa modulation of V1. This modulation could comprise part of the neural basis of the steady “prediction” sent from higher to lower areas in the predictive coding framework (i.e., rather than the prediction error, which, in theory, is sent in a bottom-up fashion).

The current study sought to both replicate and extend this finding to further describe the circuit mechanisms which integrate this top-down modulation and give rise to DD responses, the putative prediction error signals in the oddball paradigm. It is well known that heterogeneous classes of cortical inhibitory interneurons play a crucial role in cortical circuit dynamics^13, 16^. Inhibitory interneurons dictate the gain of PYRs, modulating feature selectivity and precision in neuronal computations^17–22^. Although it has not been verified, distinct interneurons could interact to modulate V1 processing to support predictive processing, regulating local gain in accord with past and present contextual regularities^23^ – decreasing excitability among ensembles coding for predictable stimuli and indirectly increasing excitability among ensembles coding for non-predicted stimuli (i.e., deviants). Our recent study showed that local somatostatin-positive interneurons (SSTs) in V1 of mice are necessary for the generation of prediction error responses to deviant stimuli (DD) in sensory oddball paradigms^11^. Exactly how SSTs enabled DD is unknown, and it is particularly unclear given that they heavily inhibit L2/3 PYRs in V1, where DD is enriched. Another class of interneurons, the vasoactive intestinal peptide-expressing interneurons (VIPs) in V1 could also be critical for the generation of prediction errors considering their well-established mediation of top-down inputs from ACa^24, 25^ and their mutually inhibitory interactions with local SSTs^26, 27^. As mentioned above, both ACa inputs to V1 and local V1 SSTs were deemed necessary for DD in PYRs. In this paper, we sought to elucidate how VIP activity relates to ACa inputs and V1 SSTs during the oddball paradigm and, further, to test whether VIPs are also critical for DD in V1. We hypothesized that VIPs could serve as a mediator in top-down modulation of V1 during the oddball paradigm, and that this could serve to disinihibit subsets of PYRs by inhibition of SSTs, enabling DD via an ACa to VIP to SST to PYR disinhibitory circuit.

Interneurons, besides serving to spatially modulate the gain of PYRs, can also enhance the generation and maintenance of cortical oscillations^28^. The study of these oscillatory rhythms, either in intracranial local field potential (LFP) or scalp EEG recordings, affords a distinct window into the circuit dynamics that affect perception, underlie cognition, and are altered in disease^29–36^. Through the emergence of rhythmic synchronized activity in distinct bandwidths (e.g. theta, alpha, beta, gamma), neuronal networks optimize information processing and, importantly, provide temporal windows that gate or route long-range inputs, structuring their influence on ongoing local processing^37^. For example, during active attentional tasks in non-human primates, top-down or feed-back connections in the visual system exhibit activity in the high alpha or beta domain (9-25Hz), while feed forward and local connections exhibit oscillatory signatures in the delta/theta (3-8Hz) and gamma bands (25-10 Hz)^10, 38^. In primary sensory regions, DD signals appear to occupy mainly delta/theta frequencies (3-8 Hz) in both visual^11^ and auditory oddball paradigms^39^, supporting the notion that DD signals are prediction errors that are “fed-forward” in cortical networks. However, the frequency band that the top-down modulatory circuit from ACa to V1 occupies in the oddball paradigm has not been established. Guided by recent work on inter-cortical interactions in top-down and bottom-up directions in primates^40^, we hypothesized that this connection would be expected to show maximal synchrony in the alpha-beta bands.

Here we tested these frequency and cell-type specific hypotheses of predictive processing in V1 during the oddball paradigm. We first replicated the finding that V1 exhibits bona-fide DD-like responses in the LFP, which are most prominent in the lower theta-band (3-8 Hz). We then go on to show that interregional synchrony between ACa and V1 is strongest in the higher-theta/alpha-band, peaking around 10 Hz. This interaction was directional during redundant (predictable) stimuli, with larger granger causality flowing from ACa to V1 than V1 to ACa (i.e., top-down). Further, with two-photon imaging during the oddball paradigm, we show that L2/3 PYRs exhibit deviance detection in V1 (replicating past work), but VIPs, SSTs, and ACa-inputs do not, showing similar response magnitudes to both the deviant and context-neutral stimuli. On the other hand, with redundant stimuli (i.e., prior to the deviant stimulus) theVIP and SST activity patterns diverged: VIPs show sensitization (increased responses) to redundant stimuli, while SSTs show enhanced adaptation (decreased responses). Also, artificial excitation of the ACa inputs to V1 with optogenetic activation at 6 or 10-Hz (the peak synchrony during this paradigm) drove these interneuron populations differentially, with VIPs excited and SSTs inhibited, suggesting that the ACa to V1 directional synchrony we observed in the LFP experiments served to selectively modulate VIPs and SST responses to redundant stimuli. Finally, we demonstrate that chemicogenetic suppression of VIPs eliminates this ACA-V1 synchrony as well as DD. Thus, our results suggest that top-down modulation of V1 during predictive processing is frequency specific and indirectly enables prediction-error-like responses (DD) in PYRs via a VIP-to-SST disinhibitory circuit.

## Results

### Deviance detection in visual cortex occurs in low theta-band power and phase locking

We recorded local field potentials (LFP) simultaneously from frontal and visual cortices of mice while the animal observed a classic visual oddball paradigm. Two bipolar electrodes (contacts separated by <400 µm) were implanted in the anterior cingulate cortex (ACa) and in the primary visual cortex (V1). ACa was chosen because i) it is higher in the cortical hierarchy than V1, ii) it is known to send dense top-down projections to V1^41^, and ii) our past work has shown that suppressing these projections eliminates deviance detection in V1 during a visual oddball paradigm^42^. Electrode locations were confirmed postmortem as previously described^11, 42^ (Fig S1 A,B). Visual stimuli consisted of black and white, full-field square-wave gratings at approximately 0.8 cycles per degree, drifting at 2 cycles per second, each presented for 500 ms and separated from one another by 500-600 ms of medium luminance gray screen. As previously described, we presented identical stimuli in three different contexts: when the stimulus was equiprobable (p=.125), redundant (p=.875), or deviant (p=.125; Fig 1C). During recordings, the animals were awake and responsive, head-fixed on a small treadmill so that they were walk in place (Fig 1 A-C). While locomotion is known to impact V1 activity^43^, mice did not show differences in locomotion across the trial types (deviant, redundant, control), suggesting that locomotion could not likely explain differences in stimulus processing across contexts in the oddball paradigm (Fig S1 C,D), in line with past work^11, 42^.

**Figure 1:**
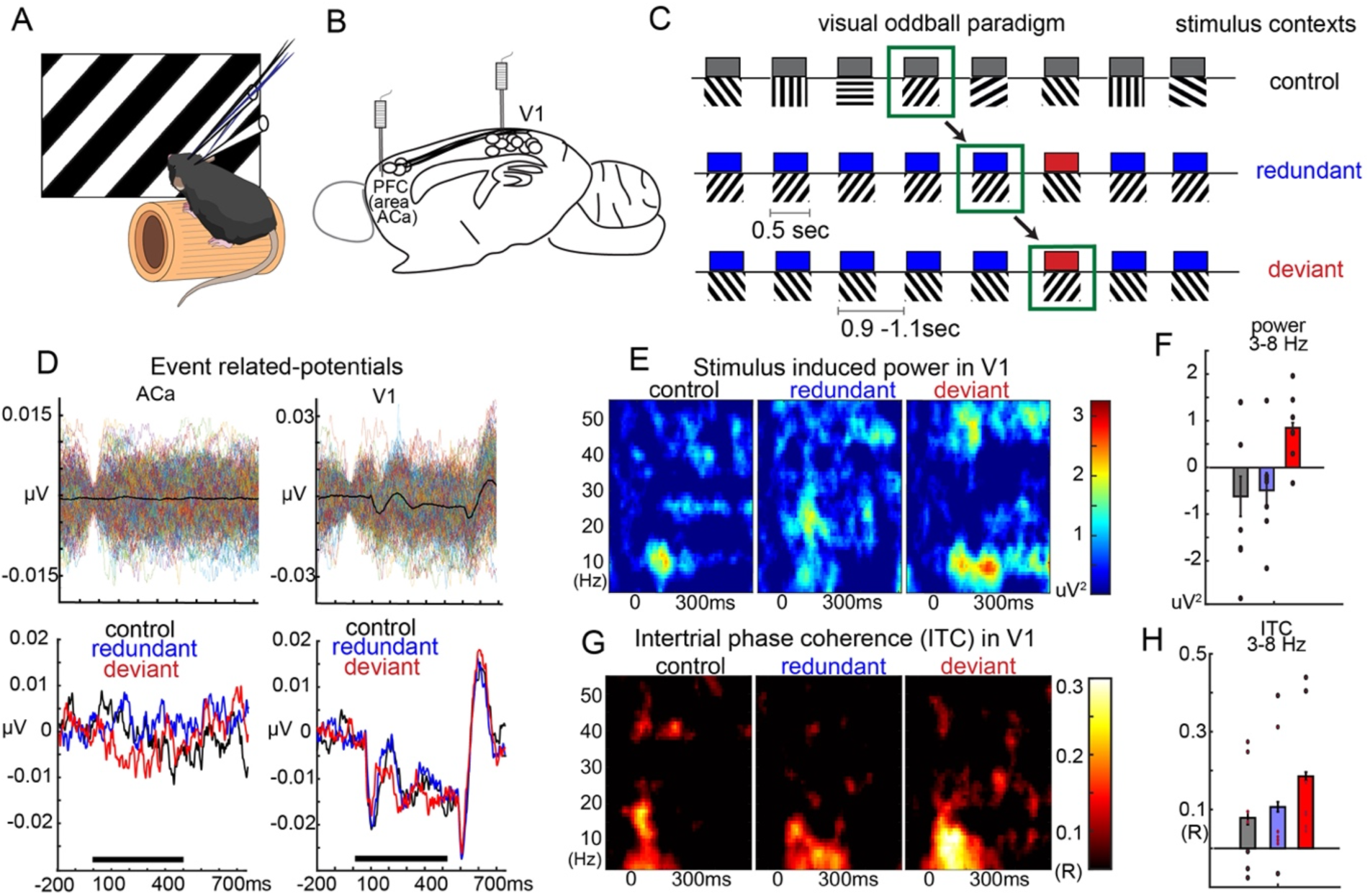
Deviance detection in V1 power and phase dynamics during a visual oddball paradigm. A) Awake mice viewed full-field visual gratings during local field potential recordings in B) V1 cortex and a prefrontal region that projects to V1: anterior cingulate area (ACa). C) Mice viewed visual stimuli in a standard oddball, oddball flip, and many-standards control paradigm. Visual responses to the same stimulus was tracked across three different contexts in which it was redundant, deviant, and neutral (equiprobable). D) (top) individual trial-activity and trial-averaged activity in ACa and V1 from a single mouse and (bottom) averaged over mice within each stimulus context show that only V1 displayed clear visually evoked responses to the stimuli. E-F) Stimulus induced power to the onset of the stimuli in control, redundant, and deviant contexts evince an increase in theta-band power to the deviant stimulus. G-H) Stimulus elicited inter-trial phase locking (ITC) to the onset of stimuli evince an early latency increase in theta-band ITC (main effect of STIMTYPE, control vs deviant; F(1,6)=10.65, p<.005).

Although many studies include an active behavioral task during sensory processing paradigms as a strategy for ensuring attention to the stimuli and engaging prefrontal regions, we specifically excluded it and overt behavior, instead employing passive paradigms, for multiple reasons. First, animals should, in theory, be able to detect unexpected stimuli even in the absence of reward anticipation or an explicit goal. Studying this function was an aim of our work. Second, recent work has shown that processing rewards or punishments activates VIP activity cortex wide^44^. This could confound our results, as we aim to study the role of VIPs in modulating sensory processing *purely with respect to sensory expectation*, which requires that any explicit rewards or punishements *not* be a part of the paradigm.

The trial averaged evoked LFP responses for a given stimulus during the control, redundant and deviant trials were generated for each mouse and averaged over mice (Fig 1D). Individual trial activity (top) and trial-averaged activity (bottom) shows that ACa does not elicit a strong visually-evoked response, while V1 does, as expected. V1 exhibited deviance detection (DD; increased activity to deviant stimulus) specifically in the lower theta band (3-8 Hz) during the oddball paradigm (Fig 1E). Stimulus-induced theta power in V1 was higher during the deviant and decreased during the redundant stimuli (Fig 1, F, main effect of STIMTYPE, control vs deviant; F(1,6)=8.66, *p<.05*). In addition, intertrial phase-locking in the low theta band in V1 was also higher for deviant trials (Fig 1, G-H, main effect of STIMTYPE, control vs deviant; F(1,6)=10.65, *p<.005*). These results are in accord with previous findings on V1 activity during context processing^11, 45^, and demonstrates that sensory processing, even in canonically “early” areas like V1, depends on context of the stimuli.

### Long-range synchrony in the theta/alpha band during the oddball paradigm suggests top-down modulation of V1

We analyzed the relationship between activity in V1 and ACa during the oddball paradigm. We probed the phase-coherence between these regions as a marker of their communication throughout the task (Fig 2A). Interregional synchrony was quantified as 1-circ variance (R-statistic) between ACa and V1 from 100 ms pre-stimulus onset to 100 ms post-stimulus offset. ACa-V1 phase-locking was statistically significant (p<.05) for frequencies between 1-22 Hz and peaked at 10 Hz (Fig 2B). Interregional phase coherence was strong and ongoing throughout the paradigm and significant even after subtracting the rest period synchrony (Fig 2B -inset). Further, this ACa-V1 coherence was strongest in layer 1 in visual cortex, consistent with this reflecting top-down inputs from ACa to V1 (Fig 2C). Further, this layer distribution is evidence against the notion that this ACa-V1 synchrony reflects simple volume conduction of hippocampal theta.

**Figure 2:**
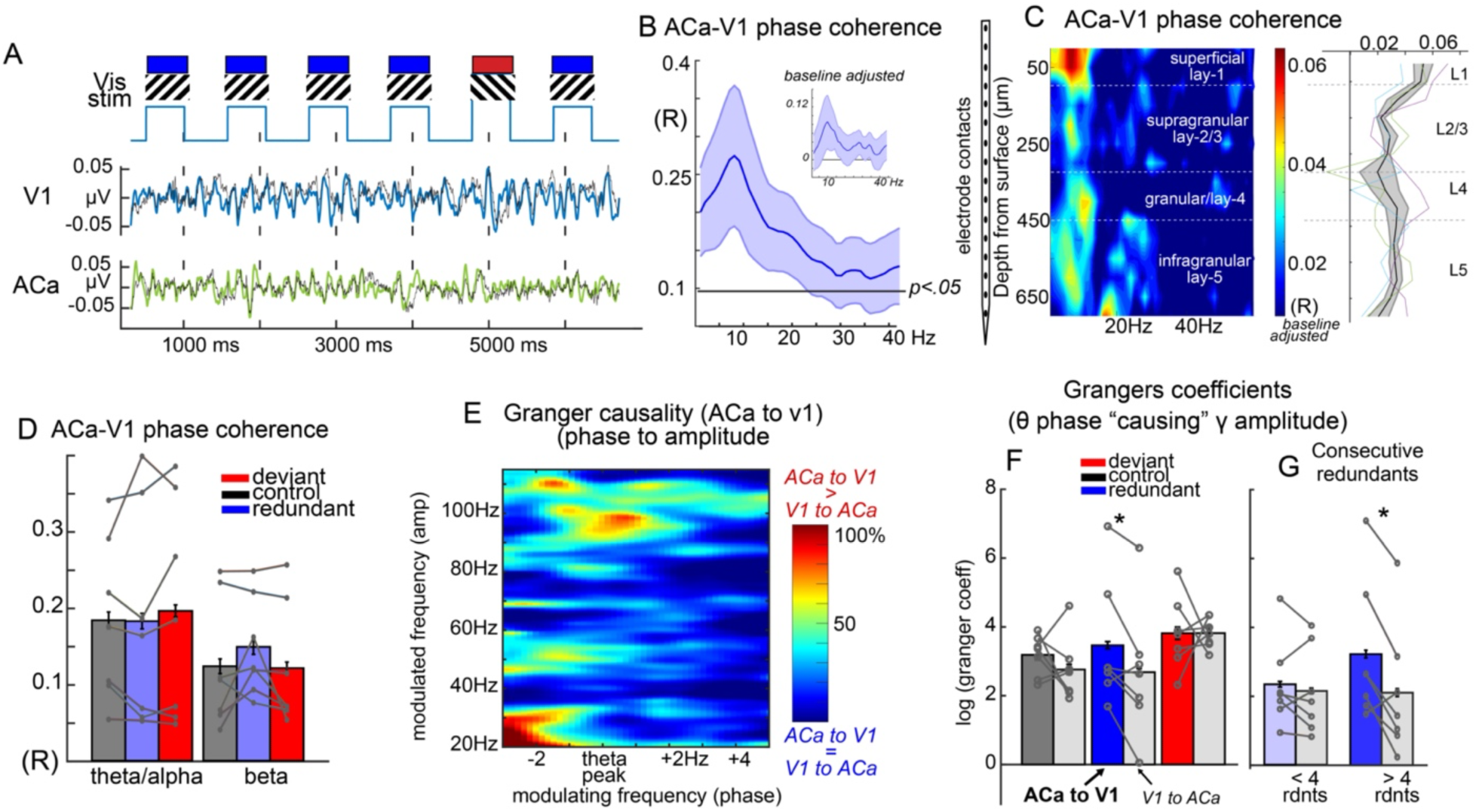
Long range fronto-visual synchrony in the theta-alpha band during the oddball paradigm. A) Bipolar electrical recordings in V1 cortex and anterior cingulate area (ACa; in mouse prefrontal cortex) displayed as ongoing unfiltered (black) and filtered local field potentials in the theta/alpha band (8-13 Hz). B) Interregional phase synchrony quantified as 1-circ variance (R-statistic) averaged across 7 mice and compared to randomized values at p<.05. Inset is subtracting pre-run (prior to the start of visual stimuli) phase synchrony. Peak is at 9Hz and extends to 20Hz. C) Recordings of a single ACa electrode combined with a multielectrode probe (16-channels) in n=3 mice, showing that ACa-V1 theta/alpha synchrony is strongest in layer 1, where ACa-axons terminate in V1. D) Magnitude of interregional phase synchrony did not differ as a function of stimulus type in theta/alpha (8-13 Hz) or beta (15-25 Hz) bands. (E) Granger coefficients of how phase of peak theta in one brain region predicted gamma power in the other region, indicating a power peak in the high-gamma frequency range, centered around peak-theta phase. F) Bar plots and individual mouse (points) data from the peak region in (D) quantified for control, redundant, and deviant stimulus conditions evince a stimulus type by direction interaction (F(3, 24)=10.0, p<.01). Only the redundant condition exhibited significantly stronger top-down vs bottom-up granger causation (t(6)=2.57, p<.05). (G) This directional difference was stronger after 4 redundant stimuli post deviant stimuli (t(6)=3.30, p<.05 vs t(6)=0.97, p=.36).

Interestingly, the magnitude of this coherence did not differ as a function of stimulus context (Fig 2D), bringing into question how this apparent synchrony relates to predictive processing during the oddball paradigm. One possibility is that this interregional coherence reflects a bidirectional modulation, with top-down inputs conveying “predictions” and bottom-up outputs conveying “prediction errors”, and the relative strength or influence varying from bottom-up to top-down depending on how the current stimulus fits with the internal model of the environment (i.e. which stimuli are and are not likely). To test this, we carried out granger causality analysis to determine how theta-phase in one region predicted the gamma power in the other at 1/8^th^ of a theta cycle in the future (similar to other work in cortico-cortical circuits^38^). Previous research has demonstrated that high-gamma power in neocortex reflects synchronous local neuronal activity^46^ (and also correlates strongly and directly with the fMRI BOLD signal^28, 47^). Granger coefficients were quantified bidirectionally (ACa to V1 [“top-down”] and V1 to ACa [“bottom-up”]) and scaled as percentage of top-down minus bottom up divided by average of both, from 100ms pre to 100ms post-stimulus. In general, ACa theta granger-caused V1 gamma power at much higher levels than the reverse direction, and the strongest ACa to V1 causation for each recording’s “peak” theta (ranging from 4 to 12 Hz) was in the high gamma-power (80-120 Hz; Fig 2E). Bar plots and individual mouse (points) data from the peak region in Fig 2E quantified for control, redundant, and deviant stimulus conditions, show a stimulus-type by direction interaction (F(3, 24)=10.0, p<.01). The phase of PFC-theta significantly Granger caused V1-gamma during redundant trials (Fig 2F, t(6)=2.57, p<.05), and this effect is enhanced for later redundants in the sequence (Fig 2G, t(6)=3.30, p<.05 vs t(6)=0.97, p=.36), suggesting that it scales with how well the current stimulus matches predictions (which are presumably built on accumulated evidence from preceding trials^2^).

To summarize, ACa and V1 synchronize in the theta/alpha band. During the most predictable stimuli, the direction of theta/alpha modulation is strongest from ACa to V1 (rather than V1 to ACa). That is, when the bottom-up information about the visual stimulus, in V1 is most consistent with the contextual regularities (i.e. redundant), the ACa-to-V1 influence is maximal. This suggests that ACa inputs convey predictive information about current stimuli to V1, consistent with past theories.

### Heterogenous cell-type and top-down dynamics in ACa-V1 circuits during the oddball paradigm

We next investigated how ACa projections to V1, as well as how specific layer 2/3 V1 neurons (VIPs, SSTs, and PYRs), are active during the oddball paradigm to further test this model of top-down predictive processing from ACa to V1. Our past work showed that while layer 2/3 PYRs in V1 show deviance detection (DD; i.e. increased responses to deviant stimuli relative to controls) and stimulus specific adaption (SSA; i.e. reduced responses to redundant stimuli relative to controls), the ACa terminals in layer 1 (L1) of V1 show *neither* DD nor SSA^42^. In line with our LFP results (Fig 1D, 2F), this suggests that these top-down inputs are not responding to the current stimulus, but, perhaps instead, could be sending information about the currently “predicted” stimulus.

Our past work shows that SSTs are necessary for V1 DD^11^, but it remains unclear exactly what role SSTs play during the oddball paradigm. In theory, SSTs could i) passively mediate a tonic disinhibition (of L2/3 PYRs which exhibit DD) during the highly predictable stimulus train (and thus immediately prior to the deviant stimulus) or ii) carry out a more “active” disinhibition, selectively releasing DD-exhibiting PYRs from inhibition from other interneurons by directly inhibiting other interneurons the presentation of a deviant stimulus. Recording SSTs during the oddball paradigm would help clarify their role. If the mechanism is passive (i), they should show increased adaptation during redundant phase, indirectly disinhibiting the PYRs they target. If their role is active (ii), they should show increased activity during the deviant stimulus (i.e., they should show a form of DD as well), temporally prior or concurrent with PYRs exhibiting DD to the deviant stimulus.

Fast two-photon microscopy (28 Hz resonant scanning) was employed to record neuronal activity from cortical layers 1-2/3 in V1 (50-300 µm below the brain surface) as awake, head-fixed mice viewed the oddball and many-standards control sequences (described above). We imaged the activity of layer 2/3 PYRs (n=4 mice, 323 neurons), SSTs (n=8 mice, 328 neurons), VIPs (n=9 mice, 284 neurons), and ACa axons in V1 (n=4 mice, 100 synaptic boutons and axonal segments which were deemed functionally uncorrelated [see methods]). Transgenic mice expressing cre-dependent GCaMP6s crossed with tm1.1-VGluT-, SST- and VIP-cre-lines were used (Fig 3A-C) except in experiments recording ACa axons in V1, where AAV1-Syn/Cag-GCaMP6s was virally expressed in ACa (Fig 3D) and imaged in L1 of V1 as previously described^42^. Visual stimulation was the same as reported in Fig 1 and 2. For analysis, only “visually responsive” neurons or boutons/axon-segments (showing >1 standard deviation from average response over baseline to at least one stimulus [out of 4 orientations: 0, 90, 45, 135 deg] under at least one condition [control, redundant, deviant]) were considered (89% of neurons recorded for PYRs; 98% for SSTs; 93% for VIPs; 89% for ACa axons). If a neuron/axonal-segment showed significant activity to more than one oriented stimulus, we plotted and used only one (the strongest after averaging across contexts) for statistical tests (Fig 3 A-D)^14^.

**Figure 3.**
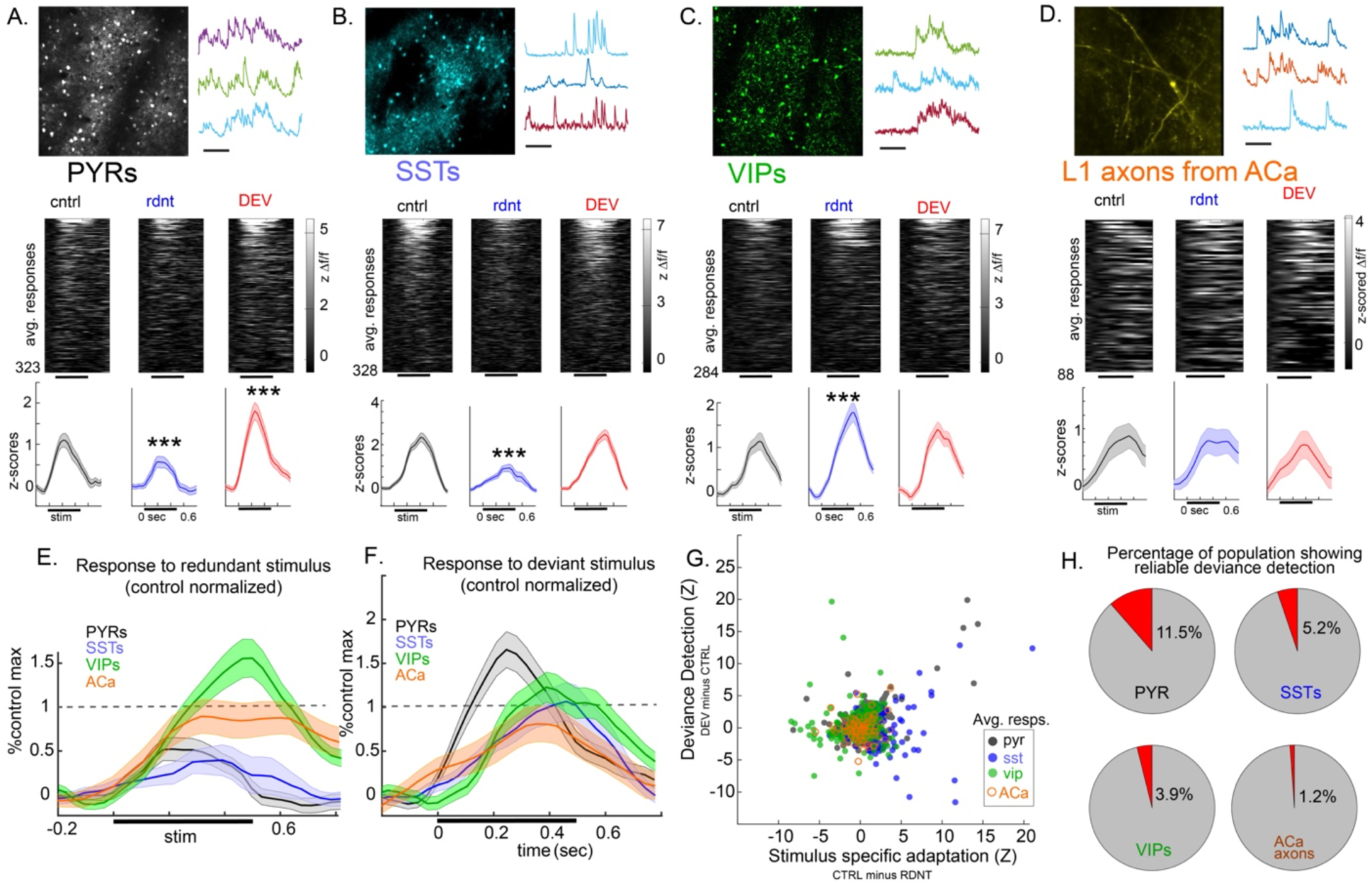
Cell-type specific dynamics in the V1-ACa circuit during the oddball paradigm. A. Two-photon calcium imaging carried out in pyramidal neurons (PYRs; n=4 mice, scale bar= 15 sec) during a visual oddball paradigm; (middle) rasterplot of average responses to the same stimulus when it is the control, the redundant (4^th^ in sequence), and the deviant stimulus in context (over 10 trials for each). Plotted are 323 “visually responsive” neurons; (bottom) averages across neurons show significant stimulus specific adaptation (redundant vs control) and significant deviance detection (deviant vs control). B) Same as A but for somatostatin positive neurons (SSTs; n=8 mice; 328 neurons, 98% of recorded); (bottom) averages across SSTs show significant stimulus specific adaptation (redundant vs control) but not deviance detection (deviant vs control). C) Same as A, but for vasoactive intestinal peptidergic neurons (VIPs; n=9 mice; 284 neurons, 93% of recorded); (bottom) averages across VIPS show significant *INVERSE* stimulus specific adaptation (redundant vs control) but not deviance detection (deviant vs control). D) Same as A, but for axon segments and boutons (regions of interest; ROIs) in layer 1 of V1 from putative ACa projection neurons (ACa; n=4 mice; 89 neurons, 89% of recorded); (bottom) averages across ACa ROIs show neither significant stimulus specific adaptation (redundant vs control) nor significant deviance detection (deviant vs control). E) Average responses to the redundant stimulus for each cell/ROI scaled relative to the maximum of the average response to the control stimulus within each cell/ROI-type. F) same as (E) but to the deviant stimulus. G) Scatterplot showing computed “DD” and “SSA” for each neuron. Different colors represent cell/ROI types. H) Percentage of each cell/ROI-type showing “true deviance detection”: i.e. response to deviant is largest and is more than 1.67 standard deviations larger than response to control stimulus in two separate sets of trials (5 even vs 5 odd trials). The estimate for PYRs is consistent with our past work^42^. ***p<.005

As previously reported^42^, the PYRs showed DD (responses to deviant vs control: paired t(320)=3.17, p<.005), and SSA (redundant vs control; paired t(320)=-3.5,p<.001) (Fig 3A,E-F). Interestingly, SSTs did not show DD (paired t(326)=0.53, p=0.60), but showed strong SSA (Fig 3B,E-F, paired t(326)=-6.95,p<.001). VIPs also did not show DD (Fig 3C, deviant vs control: paired t(282)=1.2, p=0.233) but, surprisingly, were more during active to redundant stimuli (redundant vs control; paired t(282)=2.82,p<.005), suggesting that VIPs exhibit a form of repetition sensitization or reverse adaptation. Consistent with our past work^42^, stimulus responsive ACa axons showed neither DD or SSA. That is, ACa axons in V1 showed stimulus induced activity which did not vary as a function of stimulus-type (Fig 3D-F redundant vs control; paired t(87)=-0.17, p=.87, deviant vs control: paired t(87)=-0.56, p=0.58). A closer analysis of ACa axon activity, however, evinced greater variance across the population of inputs during the oddball paradigm compared to the during the many-standards control (Fig S2H,I; Bartlett’s test statistic^ctrl_vs_rdnt^=4.60, p<.05; BTS^ctrl_vs_dev^=11.6, p<.001; BTS^dev_vs_rdnt^=1.70, p=.20). This suggests a wider spread of input magnitudes, including more highly active and more silent inputs during the oddball, when the putative predictive model is simpler (one orientation expected -- i.e. more precise priors). During the control paradigm, when the putative predictive model is more general (8 possible orientations), there was a more gaussian spread of ACa-V1 input magnitudes across the population of axons.

Only PYRs showed marked DD. Interestingly though, DD was not present in all PYRs. We estimated that approximately 11.5% of PYRs showed reliable DD across recordings (by splitting early and later trials; consistent with our past estimate^42^). This heterogeneity of DD responses across PYRs was not simply explained by differences in stimulus feature selectivity, as PYRs selective for any orientation (O.S.I.>0.20) showed deviance detection to both their preferred orientation and to their non-preferred orientations, and non-selective PYRs also showed DD (Fig S2A,E). However, the magnitude of the deviant vs control difference, (Fig S2A) and the degree to which a given PYR showed DD (Fig S2E) was positively related to the cell’s orientation selectivity, suggesting that DD may reflect a non-specific gain modulation in V1.

ACa axons, SSTs, and VIPs did not differ in their activations to deviant stimuli (relative to control; Fig 3F). These cell types did, however, differ in their activations to redundant stimuli (either in magnitude for VIPs and SSTs, or in their directional synchrony, as for ACa). It is therefore possible that the causal role that SSTs and ACa-inputs play in supporting DD (shown previously^11, 14^) involve activity during the redundant stimulus train, prior to the deviant stimulus. Specifically, the fact that SSTs in our data do not show strong responses to the deviant stimulus (i.e., no DD) suggests that SST’s role in DD is not one of active disinhibition of PYRs *during* the deviant stimulus (hypothesis (ii) above), but, rather, by strongly adapting during the redundant stimulus, one of indirect disinhibition of PYRs in the buildup to the deviant, contextually unexpected stimulus (hypothesis (i)). Conversely, VIP neurons showed augmented responses to redundant stimuli (Fig 3E). Given known mutual inhibition between VIPs and SSTs, this sensitization of VIPs may essentially give rise to the SST inhibition, or vice versa, or both.

Interestingly, ACa inputs to V1 did not reduce the overall magnitude of their activity during the redundant stimuli either (i.e., no SSA), but, instead, exerted a stronger causal influence V1 during redundant stimuli (when predictions match sensory data; Fig 2F,G). Further, ACa inputs appeared to convey different information to V1 during mostly-predictable vs less-predictable contexts (i.e. different population variance in oddball vs control runs; Fig S2H,I), but not during predicted vs unexpected stimuli *per se* (redundants vs deviants), suggesting that ACa inputs to V1 convey *a priori* “predictions” about the current sensory data to V1. It is possible that this top-down input from ACa is modulatory and serves to amplify VIP neuron responses to predictable stimuli. This advantage that VIPs could have over SSTs would cause them to “win out” in highly predictable contexts, effectively disinhibiting subsets of PYRs during the redundant phase of the paradigm, priming them for DD responses to an unexpected stimulus. On the whole this suggests the presence of a disynaptic inhibitory circuit of ACa-to-VIPs-to-SSTs explaining the causal role of ACa and SSTs in V1 DD^42^ (Fig S6).

### Top-down drive of V1 from ACa in the theta/alpha band activates VIPs and suppresses SSTs

To further test this putative ACa modulation of VIP-SST circuits, we optogenetically activated ACa inputs to V1 during rest. We drove these inputs at a range of frequencies informed by our LFP experiments. Based on our two-photon results (Fig 3), we predicted that that driving these axons at frequencies relevant to this circuit and paradigm (see Fig 2) – namely theta/alpha frequencies –should increase activity of V1 VIPs and PYRs while decreasing activity of V1 SSTs. An AAV transducing an excitatory channelrhodopsin (ChR2) under the CaMKIIa promoter (AAV9-CaMKIIa-hChR2(H134R)-mCherry) was injected into ACa (Fig S3A,B) of mice expressing GCaMP6S in PYR, VIP, or SST interneurons. Rhythmic wide-field optogenetic stimulation was performed through a craniotomy placed at the mouse V1. Activity of PYRs, VIPs, or SSTs was imaged in V1 while 1 second bursts of 473 nm light illuminated the imaging column to activate ACa axons at 2-, 6-, 10-, 20-, or 40-Hz (20% duty cycle, squarewave pulses; .5mm radius; 12 mW per mm^2^), a power-normalized “weak” block stimulation, and a full-power block stimulation (Fig 4, A-B).

**Figure 4.**
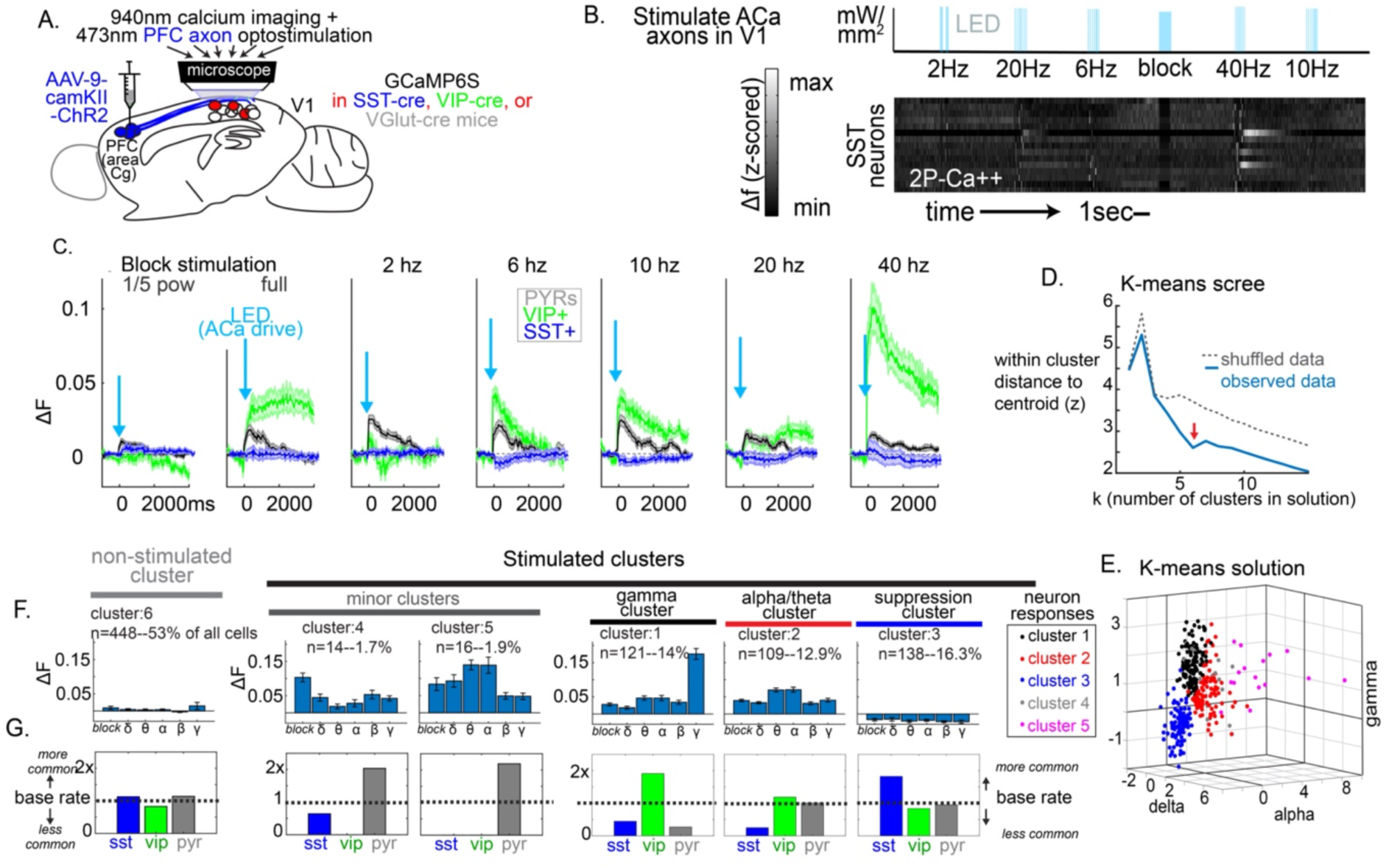
V1 activation by ACa is frequency and cell-type specific. A) a cre-dependent AAV tranducing excitatory ChR2 was injected into ACa (a prefrontal area projecting to V1) of mice expressing GCaMP6S in PYR, VIP, or SST interneurons. Activity of PYRs, VIPs, or SSTs was imaged in V1 while B) 1 second bursts of 473 nm light illumated the imaging column to activate ACa axons at 2-, 6-, 10-, 20-, or 40-Hz, a power-normalized “weak” block stimulation, and a full-power block stimulation. C). The activity of V1 cells during frequency-specific driving of ACa-inputs differed among VIPs, SSTs, and PYRs (F(10,4035)=9.60, p<.001). During 6 and 10-Hz stimulation (theta and alpha), SSTs and PYRs differed from VIP interneurons, with VIPs increasing and SSTs decreasing below baseline (all p<.001). During gamma stimulation, SSTs and PYRs also differed from VIPs, with VIPs showing dramatic responses on average and PYRs and SSTs showing moderate positive responses (all p<.001). D) Cell responses for all stimulation conditions (except weak) were standardized and had their dimensions reduced to 5 (PCA); a 1000 k-means analyses was carried out on real and shuffled data (shuffled within cells, across PC dimensions). At each iteration, we quantified the median “within cluster distance from centroids” to determine the quality of the clustering solution. Our data supports the presence of 6 stimulation clusters. E) We plotted this as a scatterplot for delta (2 Hz), alpha (10 Hz), and gamma (40 Hz), excluding the cluster with no strong ACa-drive (about half of the cells). F) We took the average centroid locations across these 6 clusters and computed their average responses to each ACa-stimulation condition. This included a non-stimulated cluster and 5 stimulation clusters. All 5 stimulation clusters showed significantly different activity to 10 Hz drive compared to block (non-oscillatory) stimulation (p<.01) except the suppression cluster. G) We plotted for each cluster the proportion of SSTs, VIPs, and PYRs relative to the overall proportion of these cells in the overall dataset. The three major stimulation clusters all comprised greater than 12% of imaged neurons each, and showed marked differences in proportion across SSTs, VIPs, and PYRs, with a gamma stimulation cluster including VIPs (which also showed significant alpha-drive) and a suppression cluster including mostly SSTs.

The activity of V1 cells during frequency-specific driving of ACa-inputs significantly differed among VIPs, SSTs, and PYRs (Fig 4C, F(10,4035)=9.60, p<.001). Follow-up analyses showed that during 6 and 10-Hz stimulation, PYR and VIP responses differed from SST interneurons, with VIPs and PYRs increasing their activation while SST decreasing below baseline (all p<.001). Consistent with our hypothesis, the cell-specific responses after top-down theta/alpha activation (6- and 10-Hz) point to a possible VIP to SST to PYR top-down disynaptic disinhibitory motif that is preferably engaged through ACa-to-V1 modulation during the oddball paradigm. This motif can be found in other areas of the cortex^18, 48^, although some direct activation of PYRs during our optogenetic stimulation could result from direct synapses of ACa on PYRs^24^.

It was also apparent that 40-Hz stimulation strongly activated VIPs. Past work on ACa-V1 circuitry showed that top-down gamma-band drive of ACa projections to V1 promotes post-error performance during an active visual attention task^25^. As we did not observe strong ACa-V1 gamma synchrony during our task-free paradigm, it is possible that the theta/alpha recruitment of V1 and the gamma recruitment of V1 represent separate mechanisms. Indeed, the heterogeneity of VIP responses to ACa drive supported this interpretation. Examining the responses of individual SSTs, VIPs, and PYRs to top-down drive, within-group heterogeneity across frequencies is apparent (Fig S4). To determine whether there were specific ensembles of cells, reaching across cell-classes, that activate to different types of top-down drive, we carried out a k-means clustering analysis on the standardized opto-evoked responses, collapsing across VIPs, SSTs, and PYRs (see Methods). Shuffling procedures and a scree-approach supports the presence of 6 stimulation clusters (Figure 4D). We took the averages centroid locations across these 6 clusters (on the non-standardized, raw data) and computed their average responses to each ACa-stimulation condition (Figure 4E). This included a non-stimulated cluster (accounting for approximately half of the V1 cells) and 5 stimulation clusters. Two of these stimulation clusters included <2% of the imaged cells (likely outliers). The remaining three major stimulation clusters all comprised greater than 12% of imaged neurons each. We plotted for each cluster the proportion of SSTs, VIPs, and PYRs relative to the overall proportion of these cells in the overall dataset. This revealed differences in proportion across SSTs, VIPs, and PYRs (Figure 4E). Clusters 1 and 2, which showed strong responses to gamma-stimulation and theta/alpha stimulation, respectively, contained very few SSTs. The former was primarily VIPs, and the latter included both VIPs and PYRs. Both clusters 1 and 2 showed significant responses to 10 Hz stimulation. Cluster 3 was a broad suppression cluster which included mostly SSTs. Interestingly, this did not show a specific effect at 10 Hz, suggesting that top-down drive inhibits SSTs regardless of frequency, but that it drives VIPs best at 10 Hz or 40 Hz.

### VIP interneurons mediate ACa modulation of V1 and support deviance detection

These results suggest that VIPs mediate the ACa to V1 10 Hz modulation (Fig 2) and are critical for DD in V1^42^. To directly test this mediating role, inhibitory cre-dependent DREADDS (AAV8-hSyn-DIO-hM4Di) were used to selectively suppress VIPs in V1 during LFP recordings during the oddball paradigm. Animals were separated in two groups: control group (no DREADDS, CNO-only) and experimental group (with DREADDS). CNO was administered intraperitoneally (IP, 5 mg/kg) to both groups after the first set of recordings. Thirty minutes after CNO administration, activity from ACa and V1 was recorded again from both groups.

In the experimental group, but not the CNO-only control, VIP suppression via CNO led to an increase in baseline power (inter-stimulus intervals) in V1 (Fig 5B, left; CNOxCONDITION interaction effect F(1,11)=5.08, p<.05), confirming a basic disinhibition of V1 by removing a source of inhibition. Consistent with our hypothesis that VIPs mediate the ACa-V1 modulation, interregional synchrony between ACa and V1 in the 6 to 12 Hz range decreased after VIP-suppression (Fig 4C, left; CNOxCONDITION interaction effect F(1,11)=5.02, p<.05). Again, DD was present in stimulus induced power in V1 in the low theta-band (3-8Hz); following chemicogenetic VIP suppression, this DD was eliminated (Fig 5 C-F; Fig S5; CNO x CONDITION x STIMTYPE interaction effect, *F(1,11)=7.10, p<.05*). Altogether these results point to VIP as an important mediator of top-down predictive processing in the visual system.

**Figure 5.**
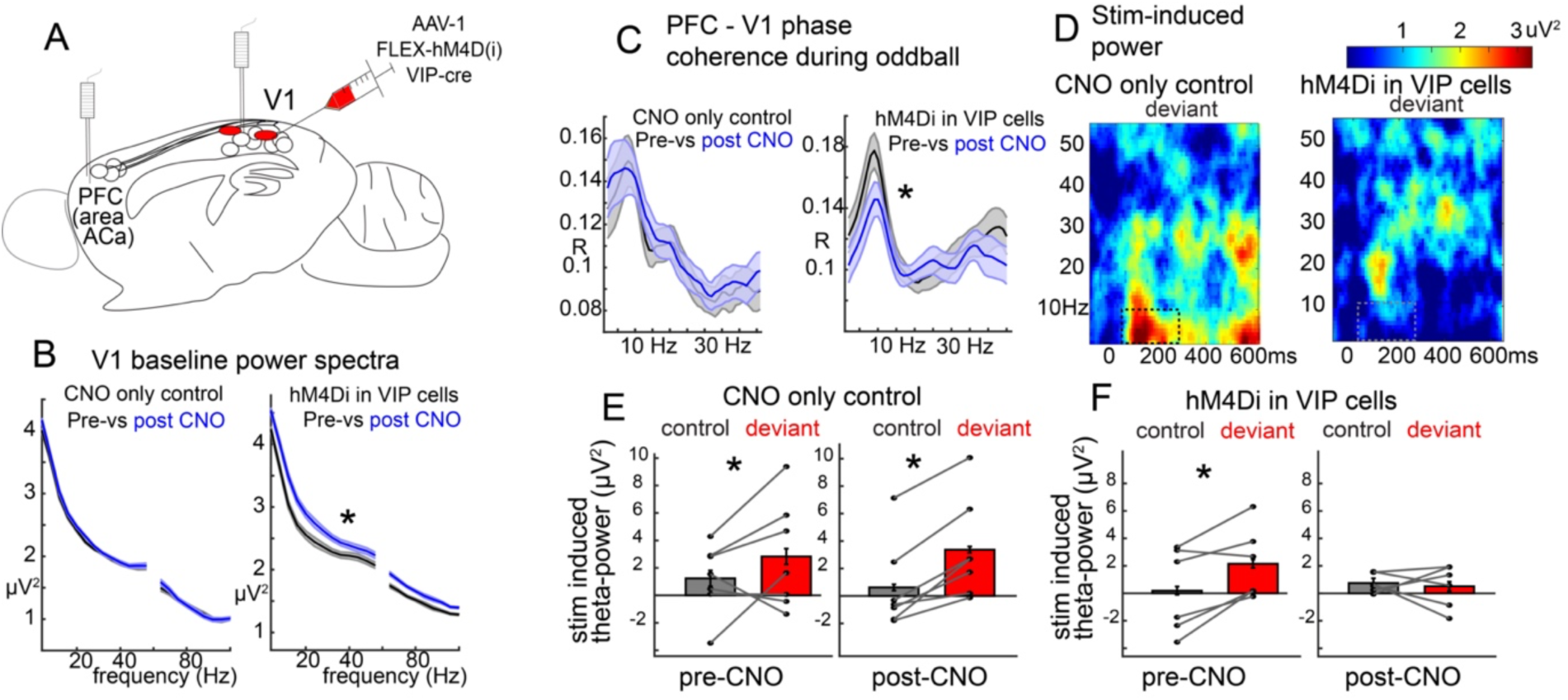
Chemicogenetic suppression of VIP-neuron activity in V1 eliminates fronto-visual synchrony and visual deviance detection. A) a cre-dependent AAV tranducing inhibitory DREADDs into VIP-positive interneurons in V1 in a VIP-cre mouse. Local field potentials were recorded in ACa and V1. B) VIP interneuron suppression increased broadband power at baseline (inter-stimulus intervals). The same effect was not seen in a CNO-only condition (left; CNOxCONDITION interaction effect F(1,11)=5.08, p<.05). C) VIP interneuron suppression decreased interregional phase-synchrony during the stimulus period of the oddball paradigm. The same effect was not seen in a CNO-only condition (left; CNOxCONDITION interaction effect F(1,11)=5.02, p<.05). D) Stimulus induced power spectra to the deviant stimuli post-CNO for the CNO-only control (left) and the VIP suppression condition (hM4D(i) in VIP interneurons; right). E,F) Stimulus induced theta-band (3-8 Hz) power from boxes in (D), points represent mice (CNOxCONDITIONxSTIMTYPE interaction effect F(1,11)=7.10, p<.05).

## Discussion

In summary, these results support a top-down circuit for contextual processing in the visual cortex (Fig S6). During a simple visual oddball paradigm, ACa and V1 are synchronous mainly in the theta/alpha band, engaging a mutually inhibitory VIP-SST circuit in V1. As both VIPs and SSTs are known to inhibit PYRs, engaging this circuit may effectively inhibit and disinhibit subsets of V1 PYRs to modulate responses to predictable stimuli while potentiating responses to future non-predicted stimuli (e.g., DD in the oddball paradigm). This model is supported by the facts i) that ACa and V1 synchronized at 10 Hz during the oddball paradigm, ii) that this synchrony showed a strong ACa to V1 directionality during highly predictable stimuli, iii) that SSTs showed strong response suppression *both* during redundant stimuli and 10 Hz stimulation of ACa inputs, while iv) VIPs, in contrast, showed *response facilitation* during redundant stimuli and 10-Hz stimulation of ACa inputs, and v) that suppressing VIPs disrupted ACa-V1 synchrony and V1 deviance detection. We discuss how these results fit within a predictive processing framework below.

### Top-down modulation influences local cell activity through a distinct frequency channel

The predictive coding framework posits a spatially distributed hierarchical network in the cortex involving feedforward and feedback connections which continuously modulate sensory processing (in lower areas) and prediction (in higher areas)^2, 49^. This requires that information be integrated within local circuits and across distal regions. While the spatial organization of physical connections within and between different brain regions certainly play a role, the temporal dynamics within these connections may serve to further segregate or route interregional signals to activate or suppress specific ensembles ^40, 50^.

Our results suggest that, during a highly predictable sequence of visual stimuli, this can occur through ACa and V1 phase-locking in the theta/alpha band. Interestingly, this synchrony may serve to potentiate responses in specific neural populations in V1 to elicit different responses at the cell-specific level. Our optogenetic driving experiments support this notion, showing that ACa inputs to V1 driven at 6- or 10 Hz elicit distinct and opposite responses in interneuron populations (SST and VIPs). This same pattern of increased VIP activity and decreased SST activity was observed during the redundant phase of the oddball paradigm, when top-down theta/alpha influence was strongest, suggesting that theta/alpha band is critical for engaging the VIP-SST mutually antagonistic circuit necessary for visual spatial^13^ and temporal (Fig 5, and ^11^) context processing in V1.

Past work in this circuit has shown that ACa may naturally send beta-band signals during a behavioral error^25^, and that ACa-to-V1 projections target multiple interneuron types to mediate disynaptic inhibition and support visual attention^24^. Our study complements this by identifying a theta/alpha-band oscillation during non-goal-directed visual processing and its effect on V1 interneuron types. The precise frequency ranges involved in feedback and feedforward signals may nevertheless differ between specific regions and animal models. As previously mentioned, in directed attention tasks with non-human primates (NHPs), feedback activity in the visual system is driven at the alpha/beta bands, while theta and gamma bands are associated with feedforward signals^40, 51^. It is possible, however, that our theta/alpha range (6-12 Hz) could be functionally similar to this studies alpha/beta range (10-20 Hz). The exact discrepancy could be due to simple differences in brain size between mice and NHPs, or due to the nature of the task itself (active vs passive paradigms; as directed tasks involve a larger information load that require multi-region integration) or due to the exact brain regions chose (as past work shows slightly varying bandwidths for the “top-down beta” modulation across cortical regions^40, 51^). Nevertheless, our results are consistent with the core idea of frequency channels being a routing mechanism for feedback modulation and feedforward information^40^, with new evidence provided here that these temporal dynamics serve to engage specific cell populations.

### VIPs modulate prediction in the absence of active report

We found that VIP interneurons are more active during predictable stimuli, suggesting response facilitation during the redundant phases of the oddball paradigm. VIPs did not exhibit a DD response (i.e., they are not differentially more active to a deviant stimulus compared to a control/neutral stimulus; figure 3). Further, chemicogenetic inhibition of VIPs both altered interregional synchrony between prefrontal and visual cortices and altered context-dependent modulation of V1 responses to visual stimuli (i.e., DD; figure 5). Thus, we propose that VIP activity, through top-down modulation, mediates the ability of V1 to generate differential responses to predictable-vs-unexpected stimuli (i.e., deviance detection), not just by direct inhibition of PYRs coding for the predictable stimuli, but by an indirect disinhibition of PYRs selective for stimuli other than the predictable stimulus (Fig S2). In other words, VIPs distribute the “predictions” of a sensory context through a pattern of inhibition and disinhibition. In figure S6, we have provided a schematic based on current and past observations (with some testable assumptions about connectivity) to facilitate future work into deviance detection circuits. Future work is necessary to more comprehensively test this ACa^long-range^-VIP^V^^1^-SST^V^^1^-PYR^V^^1^ deviance detection circuit, including studies which selectively inhibit each element while imaging the other, during different phases of the paradigm. Results here make significant strides to formulating a working model of this deviance detection circuit (Fig 6). Further, more work is needed to better understand the heterogeneity present in adaptation vs facilitation vs deviance detection in VIP, SST, and PYR populations (Fig 3G,H and Fig S2), as well as the roles of other cell types within V1, including parvalbumin positive interneurons (PVs) and neuroglioform cells in layer 1.

These findings and interpretations are largely consistent with past work in VIPs in neocortex. During locomotion, VIP interneurons are responsible for enhancing responses of weak sensory signals through the same VIP-SST disinhibitory motif that allows for a subset of cells to be highly excitable, responding promptly and strongly to stimuli^52^. Importantly, we did not find that mice showed any differential locomotion to deviant vs control vs redundant stimuli, so the modulatory role of VIPs in our paradigm cannot be simply explained by this function. In general, VIP+ interneurons are known to modulate neural activity throughout the cortex across brain-states and behaviors via a number of circuit motifs^53^. For example, VIPs play a disinhibitory role in modulating PYR neuron activity by inhibiting other interneuron types, PV and SST neurons^48, 54^. The fact that VIPs are localized in superficial cortical areas, which receive the majority of top-down inputs from higher regions, suggests that they may mediate a form of disynaptic inhibition and disynaptic dis-inhibition of multiple cell types^55^. Evidence for such motifs have been found in multiple regions of the mouse cortex^26, 43, 48, 54–56^, and serve to elicit differential responses in brain networks depending on the task and context.

Much of the past research done on the role of cortical VIPs during visual novelty processing has involved mice and the presence of a trained behavior output, often conditioned through reward and punishment, and involving locomotion as a part of the behavior^43, 44, 52, 57^. In this study, we sought to understand the functional dynamics of VIP interneurons in a passive oddball paradigm in the absence of an active report and an anticipated reward. Notwithstanding the fact that passive, untrained, or non-goal driven behavior makes up a large part of any animal’s natural life, the value of a strickly passive oddball paradigm is clear from the clinical neurophysiology literature. One of the best replicated biomarkers of schizophrenia – the mismatch negativity (an EEG index of DD) – involves a purely passive sensory sequence (i.e. the oddball sequence)^15, 58, 59^. Interestingly, our finding that VIPs are more strongly active to predictable – but not unexpected– stimuli contrasts some findings in which VIPs were shown to have higher activation to novel images in comparison to familiar ones ^57^. A key difference here may be the nature of the paradigm; Garret et al (2020) used a go/no-go task in which the mice had to report the changes in their visual perception by licking, and by doing so, receiving a reward. This task involves both locomotion and reward, and recent work has shown that both modulate cortical activity. In particular, reward and punishment lead to VIP activation^44, 57^. Therefore, it is unclear whether past findings of VIP activation to novel stimuli are associated with the novel stimulus itself, or with the reward administration, or both.

It is possible that a wide range of VIPs functionality is a testament to its flexible and yet standard response in brain systems: VIPs are active to enhance the difference between neuronal populations engaged in processing different inputs, regardless of the main driver of VIP activity. Distal cortical and subcortical inputs modulate VIP responses to stimuli to create leverage in stimulus evoked responses that through disinhibition may generate an excitability gradient in local circuits that allow for different ensembles of cells to have a distinct response given the input. This is a circuit mechanism that allows for gain modulation in neuronal populations in accord with the needs of the task at hand (passive exploration or reward seeking).

### Top-down modulation is necessary yet not specific to redundant trials

Our past work has shown that top-down inputs from ACa to V1 are necessary for visual DD^42^, which is consistent with previously identified roles of prefrontal and/or top-down modulation from hierarchically higher areas in the cortex in supporting basic sensory mismatch processing in humans.^60, 61^ Our results here show steady activity in ACa axons in V1, time-locked to stimulus presentation, which do not show differential responses during the redundant vs control vs deviant phases of the paradigm. This is also consistent with our past findings^42^. This is potentially perplexing: how can top-down inputs contextually modulate responses to a visual stimulus in V1 if those top-down inputs themselves appear indifferent to context?

One possible explanation is that the top-down inputs are carrying predictive information that is not evoked by the current stimulus, but is signaling what internal models would hold that the current stimulus could be. When the bottom-up signals – i.e., the inputs from thalamus/layer-4– concord with the top-down predictions from ACa, responses to the current stimulus may be attenuated. Our Granger analyses of LFP data are consistent with this interpretation, as ACa-to-V1 causal influence is strongest during redundant trials (when top-down and bottom-up are in concordance). On the other hand, when bottom-up and top-down do not match, as they would during the deviant stimulus, responses to the current stimulus are enhanced, leading to most of the V1 activity to be driven locally or in the bottom-up direction. Past work in primates suggests that bottom-up processing streams occupy lower theta-band frequencies, consistent with our finding (and others’^59, 81, 82^) that theta (3-8Hz) power is strongest to the deviant stimulus. What follows from our hypothesis would be that during deviant stimuli, the direction of LFP causation should go from thalamus to V1, or from layer 4 to layer 2/3. Future work could test this with multielectrode recordings.

In further support of the interpretation that ACa axons are sending predictive information to V1 is that while the average of the magnitudes of input from ACa to V1 do not differ across contexts (oddball vs many-standards control Fig 3), the variance of activity across that population was wider during the oddball compared to the many standards control (Fig S2H,I). This suggests that activity of top-down inputs was more specific – including more clearly active and more clearly non-active ACa-V1 inputs – when the probability of the next stimulus was more certain (i.e. very likely the redundant stimulus) compared to when the next stimulus could be one of 8 orientations (i.e. during the control paradigm). In the latter case, the activity of ACa-V1 inputs appeared to aggregate around more intermediate values. Interestingly, the variance of axonal activations did not differ between the redundant and the deviant stimuli during the oddball paradigm, further suggesting ACa-V1 inputs do not convey stimulus evoked activity (Fig 1D) back to V1, but, rather, they comprise anticipatory inputs containing the predictive information.

### Clinical implications

Predictive processing and deviance detection hold significance in both basic and clinical neuroscience, as such functions are hypothesized to be disrupted in schizophrenia (SZ) and other psychotic disorders^62^. In human EEG recordings during a basic visual or auditory oddball paradigm, deviant stimuli elicit a pre-attentive mismatch negativity (MMN) event-related potential at mid-latencies (100-200ms) after the onset of a contextually deviant stimulus that, like the LFP DD signal we identify in this study, strongly correlates with increased oscillatory power in the theta band. People with SZ show reduced MMN and stimulus induced theta power to deviant stimuli^59^, and this biomarker is strongly correlated with cognitive symptoms in the disease^58, 63^, suggesting that DD dysfunction could index core information processing deficits in SZ.

SZ involves neuropathological dysfunctions of prefrontal cortex (PFC) ^64–66^, which weigh heavily in many leading theories of SZ pathophysiology^67–69^. However, neuropathology in basic sensory cortices^64, 70^ and deficits in sensory cortical processing – including auditory and visual domains – are also reliably present in the disease and are notably independent of cognitive or attentional modulation^71–73^. Such deficits could be just as crucial for explaining disease pathology and predictive processing deficits as prefrontal pathology^74^. By studying the dynamics of the homologous murine ACa-V1 circuit (ACa is a visually-projecting PFC region in mice), our results might tie together these two lines of evidence to show how erroneous sensory processing can affect -- and be affected by--dysfunction in regions higher in the cortical hierarchy like PFC, tertiary sensory regions, or parietal cortices. This prefrontal dysfunction then fails to provide the appropriate context to sensory areas, creating a disrupted loop of information processing and generating a schism between an internal model of the world and sensory inputs from the world. Specifically, our results suggest how top-down circuitry, alpha-band oscillatory disruptions, and cortical interneuron subpopulations could all contribute and relate to these core phenomenological features of the disorder.

## Acknowledgments

This work was funded by the National Eye Institute (R01EY033950, Hamm), National Institute of Mental Health (K99/R00MH115082, Hamm; F32MH125445, Ross), Brain and Behavior Research Foundation (YI30149; Hamm), and the Whitehall foundation (2019-05-443; Hamm).

## Author Contributions

JPH designed the study; GB, JTH, JMR, AMR, and CGG collected data; GB, JMR, and JPH analysed data; GB and JPH wrote the manuscript; All authors edited the manuscript; JPH and DSP conceptualized the work; JPH supervised the work; JPH, JMR, and DSP acquired funding.

## Methods

### Animals, Surgery, and Training

Experiments were carried out under the guidance and supervision of the Georgia State University (GSU) Division of Animal Resources and were approved via Institutional Animal Care and Use Committee (IACUC) at GSU. Adult C57BL/6 mice (n=47, P60 to P120, from Jackson Laboratories) were used. Transgenic lines were made using mice expressing cre-dependent GCaMP6s (tm162(tetO-GCaMP6s, CAG-tTA2)) crossed with tm1.1-(VGluT-), SST- and VIP-cre lines.

For experiments involving calcium imaging, head-plate fixation and craniotomy surgeries were carried out together as previously described^75^. A hole with diameter of 3 mm was drilled in the mouse skull in left V1 (coordinates from bregma: X =2 mm, Y = −2.92 mm), followed by the removal of the skull and exposure of brain surface; dura matter was conserved. A cover glass was placed and sealed at the hole location. Then a titanium head-plate was attached to the mouse head to allow for their fixation on the microscope. For calcium imaging or optogenetic manipulation of ACa axons, virus injections were done 2 to 3 weeks prior to head-plate fixation and craniotomies. a small hole was drilled in the mouse left ACa (coordinates from bregma: X =0.35 mm, Y = 1.98 mm, Z = 0.9 um from brain surface); a micro-syringe attached to a stereotaxic apparatus was used to inject 0.75 ul of a 1:1 solution of PBS and channelrhodopsin (pAAV9-CaMKIIa-hChR2(H134R)-mCherry) or GCaMP6s (pAAV.Syn/Cag.GCaMP6s.WPRE.SV40) over a 10 minute period (0.075 µl/min) to each mouse.

For local field potential experiments, head-plate attachment was carried out prior to electrode implantation. During the latter, two bipolar electrodes twisted together (with contacts spaced approximately 200 µm apart) were inserted 0.5 mm below the dura in stereotaxically defined ACa and V1 (coordinates from bregma: ACa, X =0.35 mm, Y = 1.98 mm; V1, X =2 mm, Y = −2.92 mm), and grounded on the skull contralateral to the target areas, totaling 4 contacts for each region (two at target, two grounded). For experiments with LFP and VIP chemicogenetic suppression, 0.75 µl of 1:1 diluted AAV8-hSyn-DIO-hM4Di-mCherry was injected in V1 starting at 0.9 mm deep and moving up to 0.5 mm from the brain surface continuously during the whole injection period (at a rate of 0.04 mm/min) to assure an even and widespread expression of the virus. Injections were done in VIP-cre mice at the same time as head-plate fixation. All animals that went through surgery were anesthetized using 3% isoflurane and received pre and post care medication appropriately (5 mg/kg carprofen, IP). Prior to recordings, mice underwent at least 3 training sessions to acclimate them to head-fixation and the visual stimuli, as previously described^76^.

### Visual Stimulation

Visual stimulation was presented on a flat TV screen at a 45° angle from the animal axis, approximately 15 cm from the eye, using Psychophysics Toolbox on MATLAB (Mathworks). Full-field, black and white, sinusoidal moving gratings were presented at 100% contrast, 0.08 cycles per degree, two cycles per second, at 8 possible orientations (30°, 45°, 60°, 90°, 120°, 135°, 150°, and 180°). Stimuli were presented for 500 ms, with an inter-stimulus interval of 500 ms of black screen. A “many standards control” (equally rare, randomly presented stimuli at all 8 possible orientations) was presented before each oddball trial to establish baseline activity. The oddball trials consisted of a repetitive sequence of one stimulus (“redundant”, either 30°, 45°, or 60° degree angles, presented 87.5% of the time), followed by a stimulus of a different orientation (“deviant”, 120°, 135°, or 180° degree angles, presented 12.5% of the time). At the latter half of the trial, the redundant stimulus is “flipped” to become the deviant, and vice versa (“oddball flip”); this way we can assess responses to the stimulus context -i.e. when in the paradigm it is shown-rather than stimulus features -i.e. what orientation it is.

### Optogenetics experiments

One second bursts of 473 nm light delivered through an LED (Bruker optogenetics module) was focused to 100 µm below V1 to activate ACa axons at 2-, 6-, 10-, 20-, or 40 Hz (20% duty cycle, squarewave pulses; .5 mm radius; 12 mW per mm^2^ at the surface), a power-normalized “weak” block stimulation (delivering the same overall power per second as the rhythmic stimulations – 2.4 mW per mm^2^), and a full-power block stimulation (1 second of 12 mW per mm^2^). These seven conditions were carried out in random order and interspersed with 9 seconds of rest between them, and each condition was repeated 10 times. No visual stimulus was shown in this run. A black tape was placed around the objective to prevent spillage of the blue light into the animal’s eyes. Virus expression and stable drive of these neurons were confirmed via histology (Fig S3A,B) and LFP recordings in ACa during V1 drive (Fig S3E-H).

### Recordings

Two-photon microscopy (28 Hz framerate; Bruker Investigator laser scanning microscope; Bruker Corporation, Billerica, MA, USA) excited by a laser (Chameleon Ultra II, Coherent Inc, Santa Clara, CA, USA) at 920nm wavelength were used to image the fluorescent calcium sensor GcAMP6s expressed in PYRS, VIPS, SST cells and ACa axons at the visual cortex of mice. The laser beam was modulated with a Pockels cell (Conoptics 350-105, with 302 RM driver) and scanned with galvometers through a water immersion objective (16X/0.80W, Nikon, Tokyo, JP). The objective lens were place on top of the animals head while a small volume of Aquasonic ultrasound gel (Parker Laboratories Inc) was placed at the site of the cranial window to bridge the objective with the imaging area and allow stability over long-duration sessions. The animals were awake, head-fixed to the microscope by their headplate, while sitting on top of a wheel free to move forward and backwards. All recordings were carried out in a dark room with the researcher present to monitor mouse wakefulness and check for signs of discomfort. Each run had a duration between 6-7 minutes. Scanning and imaging were done through Prairie View (Prairie Technologies) software (resonant galvo, downsampled to 28 frames per second, for 256×256 pixels, 3.136 µm pixel size, 802.9 x 802.9 µm field of view). A time series was recorded using Prairie View software as the mice observed visual stimuli or received opto-drive. Visual stimulus was transmitted to the monitor through an HDMI cable. The visual stimulus was converted to voltage traces and connected to the computer for stimulus recording through the Voltage Recording tool on Prairie View. Time series and stimulus voltage traces were synchronized at the onset of recording for proper alignment of neuronal activity and stimulus presentation. Optogenetic drive waveforms were converted to voltage traces and recorded at Prairie view in a similar fashion, with the signal being transmitted both to the light-stimulation driver and the computer. For PYRS, VIPs and SST recordings, images were taken 100 µm to 350 µm deep, aiming at layer 2/3 of the mouse cortex. For ACa axons, recordings took place at 50-100 µm deep, aiming at layer 1 of mouse cortex.

For the LFP experiments, the mouse was fixed to the recoding apparatus through their head-plate and free to move back and forth. Insulated cables were connected to the electrodes on the top of the head of the animal and plugged into a differential amplifier (Warner instruments, DP-304A, high-pass: 0 Hz, low-pass: 500 Hz, gain: 1K, Holliston, MA, USA). Amplified signals were passed through a 60 Hz noise cancellation machine (Digitimer, D400, Mains Noise Eliminator, Letchworth Garden City, UK), which, instead of filtering, creates an adaptive subtraction of repeating signals which avoids phase delays or other forms of waveform distortion. Multielectrode probe LFPs (Fig 2C) were recorded from a custom designed 16-channel NeuroNexus probe (750 μm length, 50 μm inter-contact distance; A1×16–3 mm 50–177; Ann Arbor, MI) inserted perpendicularly into left V1 at 100 μm/min until the dorsal-most electrode was just below the dura (deduced from real-time signals). Prior to insertion, probes were submerged in DiI dye for post-hoc anatomical validation. These LFP data were acquired from 0.1-7500 Hz upper and lower bandwidths, sampled at 10 kHz, and then low-pass filtered at 150 Hz and resampled at 1 kHz for preprocessing. Electrophysiology activity was recorded using the Prairie View software or with an Intan Recording system (multielectrode recordings). Visual stimulus timings were recorded as voltage traces at the same time as the LFPs signals for proper alignment, and the timing of these stimuli relative to LFP recording was confirmed with photodiodes placed on the video monitor. For VIP suppression, as soon as the first recordings were done, the animals were injected with CNO (IP, 5 mg/kg)^11^ and recorded again after 30 min of downtime.

### Two photon image processing

Videos were corrected for motion using the “moco” plugin on ImageJ^77^. Cellular activity was semi-manually scored using an in-house built script on MATLAB, as previously described^76^. Mean and standard deviation were calculated for all image frames and plotted for reference; rectangular sessions were manually selected around cell bodies/axon segments through a GUI in MATLAB. A PCA analysis was performed within the selected regions of interest (ROIs) to select the pixels with weights at least 80% of the maximum of the first PCA component as the final ROI, then plotted as an average fluorescence across pixels. The fluorescence time courses were displayed after each selection to verify stability across imaging experiments and healthy calcium transients. Halo subtraction was performed in the selected ROIs to exclude excess fluorescence from nearby cells. For scoring axonal/bouton ROIs, datasets were first downsampled to 9.4 frames per second to aid in scoring and detection. This is consistent with our past analysis^42^ and was necessary due to the fact that such small ROIs have faster transients and smaller signal to noise ratios. For analysis, time-courses were re-interpolated to 28 frames per second for comparison with other cell types (PYRs, VIPs, SSTs). Effectively this was equivalent to a 3 sample gaussian smooth. For axon segments and boutons, we attempted to exclude ROIs originating from the same cell (i.e. i.e. we focused those which were apparently functionally independent). That is, segment/bouton pairs showing highly similar (r>.7) activity were assumed to originate from the same cell, and thus were combined or not included (if one was less stable than the other over the imaging period).

The fluorescence traces from the resulting ROIs of soma (PYRs, VIPs, SSTs) or axons (from ACa) were converted to delta-F through a regression based smoothing approach (3-second lowess envelope)^76^. The first discrete derivative was calculated as a proxy for neuronal activity. Delta-f was z-scored within each neuron using the bottom 8% of signals across each run and then averaged across trials for each stimulus type. Only stimulus-driven cells (cells with>+1 standard deviation above pre stimulus baseline on at least one stimulus type) were considered for analysis.

For imaging cells during optostimulation, a slightly different quantification was used after the ROI extraction. Frames during which the opto-stimulus was delivered were identified by the presence of saturation artifacts in the imaging dataset. These frames and 1 frame before and after were discarded from further analyses (1.07 seconds). In contrast to the scoring of neural activity during the oddball paradigm (above), we were unable to quantify fast transient onsets in these experiments (because they started during opto-artifact). We utilized the slower decay time of the GCaMP6s calcium transients in the time period immediately following the opto-stimulation train. We quantified the average fluorescence value in the 1-second after stimulation and subtracted the eighth percentile of fluorescence in the 5 seconds occurring prior to the opto-stimulus (the baseline), and then divided the result by this baseline (deltaF/F)^78, 79^. These responses of V1 neurons were averaged across 7-10 trials within each stimulation condition.

### Local field potential signal processing and analysis

Trials with excessive signal (>≈5 std devs) in either V1 and ACa were manually excluded (between 0 and 20). All analyses were limited to the last 10 control trials and the first 10 deviants of each orientation (two per mouse). We used the 4^th^ redundant in each sequence after each deviant (also 10 total for each orientation). Analyses were combined across both orientations, as local field potential responses in mouse V1 are known to not exhibit significant orientation selectivity^80^. Trial averaged evoked responses to a given stimulus in the control, redundant, and deviant contexts were generated for each mouse and averaged over mice (Fig 1) for descriptive purposes only. Ongoing data were converted to the time-frequency domain with a modified morelet wavelet approach with 100 evenly spaced wavelets from 1 to 120H z, linearly increasing in length from 1 to 20 cycles per wavelet, applied every 10 ms from 300 ms pre- to 700 ms post stimulus onset (200ms post-stimulus offset) as previously described^11^. Stimulus induced power spectra (1-120 Hz) were computed for all three conditions (control, redundant, deviant) for each mouse and baseline corrected by subtracting the average for each frequency in the 100 ms prior to stimulus onset. Statistical analyses focused on identifying signatures of deviance detection (deviant vs control) in the low-theta band induced power and phase locking during the stimulus period, as this band has been previously shown to track deviance detection in rodents and humans^11, 59, 81, 82^.

Interregional phase synchrony was quantified by taking the phase difference for each frequency (1-40 Hz) between ACa and V1 for all measurements from 100 ms pre- to 100ms post-stimulus offset (separately for control, redundant, and deviant trials) and calculating the 1-circ variance (R-statistic) of these lags. We averaged these values compared to the random distribution at p<.05 ^83, 84^.

To determine the directionality of the peak phase synchrony between regions ACa and V1, granger causality analyses was employed to focus on how theta-phase in one region predicted high gamma-power in the other region at 1/8^th^ of a theta-cycle in the future^85^. We focused on each recording’s “peak” theta (the theta frequency with the greatest ACa-V1 synchrony – ranging from 4 to 12 Hz) and high gamma-power (80-120 Hz) as this reflects synchronous, highly local neuronal activity^28, 47^. Briefly, gamma-power in region A at timepoint “t” is treated as a criterion variable X, and is first regressed on gamma-power in region A at timepoint “t minus lag”. Then, theta-phase in region B at timepoint “t minus lag” is added to the model, and the change in R^2^ is quantified as an F-value – the so-called “Granger coefficient”. These “Granger coefficients” were log-transformed and quantified for both directions (ACa to V1 [“top-down”] and V1 to ACa [“bottom-up”]) for all timepoints and trials from 100 pre-stimulus to 100ms post stimulus offset for all mice, and averaged within conditions (control, redundant, deviant).

### Calcium imaging data analysis

For analyzing cell-level responses during the oddball paradigm, calcium imaging analyses were limited to the 10 trials in each condition (control, redundant, deviant) as described above (due to locomotion and field-of-view drift artifacts). We generated stimulus triggered averages for each neuron recorded for each of the three conditions. If a given neuron responded greater than 1.67 standard deviations above the pre-stimulus baseline activity for any one condition and any one of the orientations, we included it in subsequent analyses (between 80% and 98% of neurons imaged). Responses to only 1 oriented stimulus for each neuron (the strongest after averaging across contexts) were analyzed. We determined significant stimulus specific adaptation and deviance detection at the population level within each cell class (PYRs, VIPs, SSTs, ACa-axons) statistically as described below. Further, for each cell class, we calculated the proportion of cells showing “true deviance detection”: i.e. response to deviant is largest and is more than 1.67 standard deviations larger than response to control stimulus in two separate sets of trials (5 even vs 5 odd trials). For analyzing DD and SSA as a function of a cell’s orientation selectivity index (OSI) and selectivity relative to the orientation of the test stimulus, we quantified OSI as 1-circular variance^86, 87^ on the control trials not used for the main analysis (i.e. the 10-15 trials for each of 8 orientations prior to the final 10). Cells with greater than 0.2 OSI were considered selective. Preferred orientation for each cell was estimated as the phase of the average vector, generated by averaging all responses to the 8 different orientations.

For the optostimulation experiments, we used only runs without significant locomotion/movement artifact, including 7 to 10 trials per stimulation condition (weak, strong-block, 2-Hz, 6-Hz, 10-Hz, 20-Hz, 40-Hz). Average delta-f in the 1 second period after the LED was turned off, averaged across trials within each condition, was used for statistical analyses (see below). Each cell was an observation, and PYRs, VIPs, and SSTs were included as separate groups in a frequencyXcell-class ANOVA to determine whether significant effects of ACa stimulation frequency were present that differed across cell class.

### K-means clustering analysis of optostimulation responses

Responses of V1 neurons were averaged across 7-10 trials within each stimulation condition and combined regardless of cell class into a cells by stim stimulation condition matrix (except weak, which showed very low responses for most cells and was excluded from the k-means analysis). This 845 × 6 matrix was reduced to 845 × 5 with a principle components analysis (based on a scree-plot). We then carried out 1000 k-means analyses on real data and on shuffled data (shuffled within cells, across PC dimensions) for k=2-15 (k=number of clusters). At each iteration, we quantified the median “within cluster distance from centroids” to determine the quality of the clustering solution. We averaged these values for each value of k, creating a “scree” plot which shows how much each additional cluster adds to the solution. The point at which this curve becomes linear and parallel with the shuffled data suggests that additional clusters are no-longer necessary. This analysis suggested that our data supports the presence of 6 stimulation clusters. We took the average centroid locations across these 6 clusters (on the non-standardized, raw data) and computed their average responses to each ACa-stimulation condition. We plotted for each cluster the proportion of SSTs, VIPs, and PYRs relative to the overall proportion of these cells in the overall dataset. We specifically tested whether the 10 Hz condition differed from the block stimulation for each cluster with a paired t-test on the cells in the cluster of interest (two-tails).

### Locomotion Detection

Videos of mice during experiments were recorded at 30 fps during each experiment via a Logitech C920 HD Pro webcam mounted ≈20 cm away from the mouse’s face, illuminated by a dim 617 nm LED. Wheel motion, a surrogate of mouse locomotion, was calculated on a frame by frame basis post-hoc by singular value decomposition of manually selected ROIs using the open-source Facemap software^88^. Locomotion was binarized and analyzed similarly to LFP data, as stimulus-triggered averages C, R, and D (figure S1).

### Statistics

For LFP experiments, we carried out mixed-ANOVAs on the measures of interest at the mouse level (one measurement per mouse) with stimulus type (CONTROL, REDUNDANT, DEVIANT) and/or CNO- (PRE, POST) as within subjects factors and GROUP (hM4D(i), CONTROLS) as a between subjects factor. Significant interactions were carried out with t-tests (two-tailed). For calcium imaging experiments, we carried out repeated-measures or factorial ANOVAs on the measures of interest at the cell level with stimulus type (CONTROL, REDUNDANT, DEVIANT) or STIMULATION FREQUENCY (block, 2-, 6-, 10-, 20-, or 40-Hz) as within subjects factors. Significant effects were carried out with t-tests (two-tailed), focusing on planned contrasts between i) control and redundants (which tests for “stimulus specific adaptation”) and ii) controls and deviants (which tests for “deviance detection”) as previously described.

## Supplemental Figures

**Figure S1.**
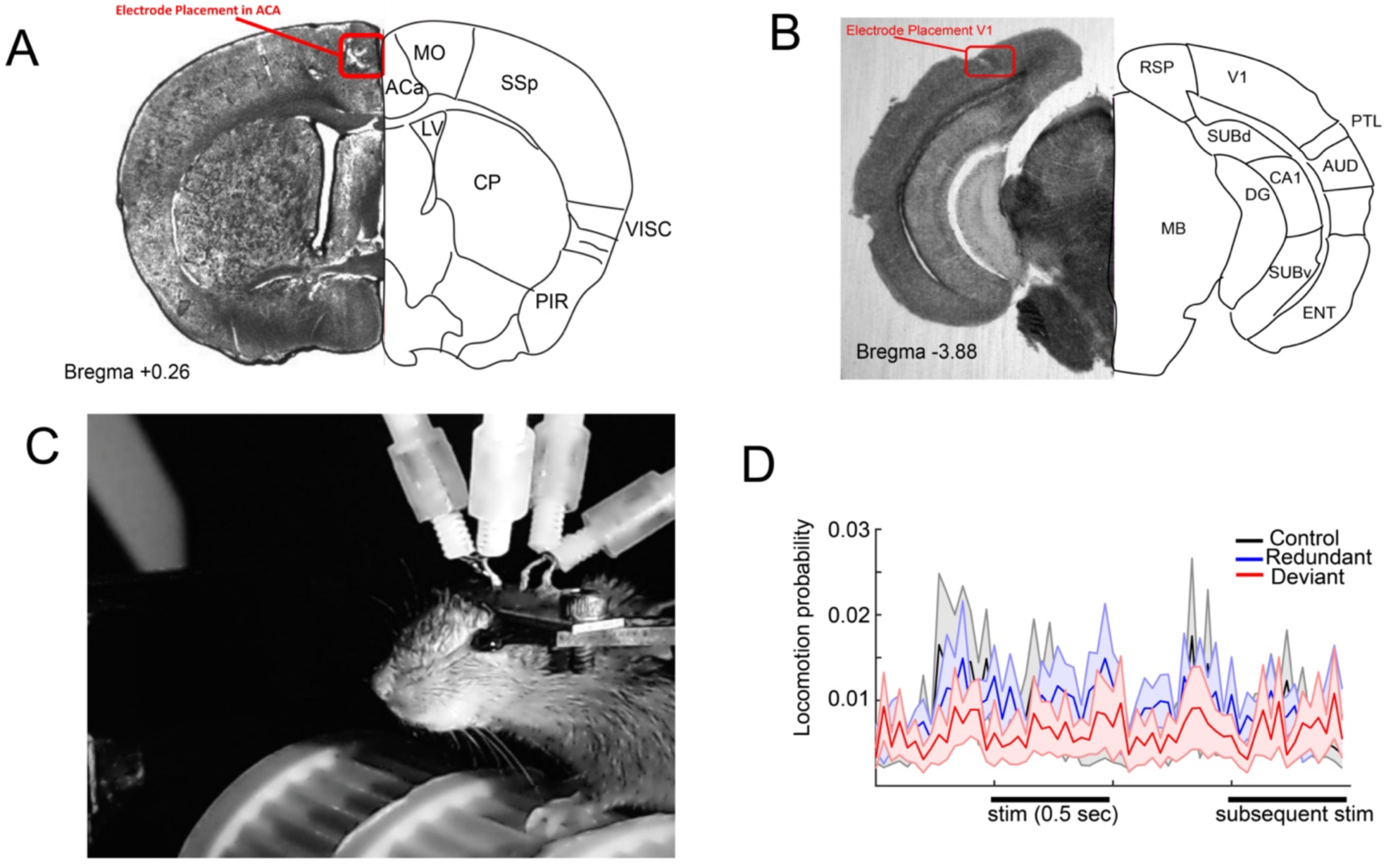
Neither deviant nor redundant stimuli evoke changes in locomotion. A) Example of anatomical confirmation of electrode placements in ACa and B) V1 was completed after recordings. C) Videos of mice during recordings were used to ensure the mouse was alert and electrodes/objectives were stable during the length of the recording, and were also used to quantify locomotion by selecting an ROI over the mouse treadmill and binarizing frames with locomotion in a subset of mice (n=14). D) Locomotion probability across trials was not different between stimulus contexts in the pre-stimulus period (F(2,12)=0.73, p=.49) or during the stimulus (F(2,12)=0.37, p=.69), nor were any t-tests between individual stimulus contexts (e.g. deviant vs control) significant in post-hoc analyses (all p>.45).

**Figure S2.**
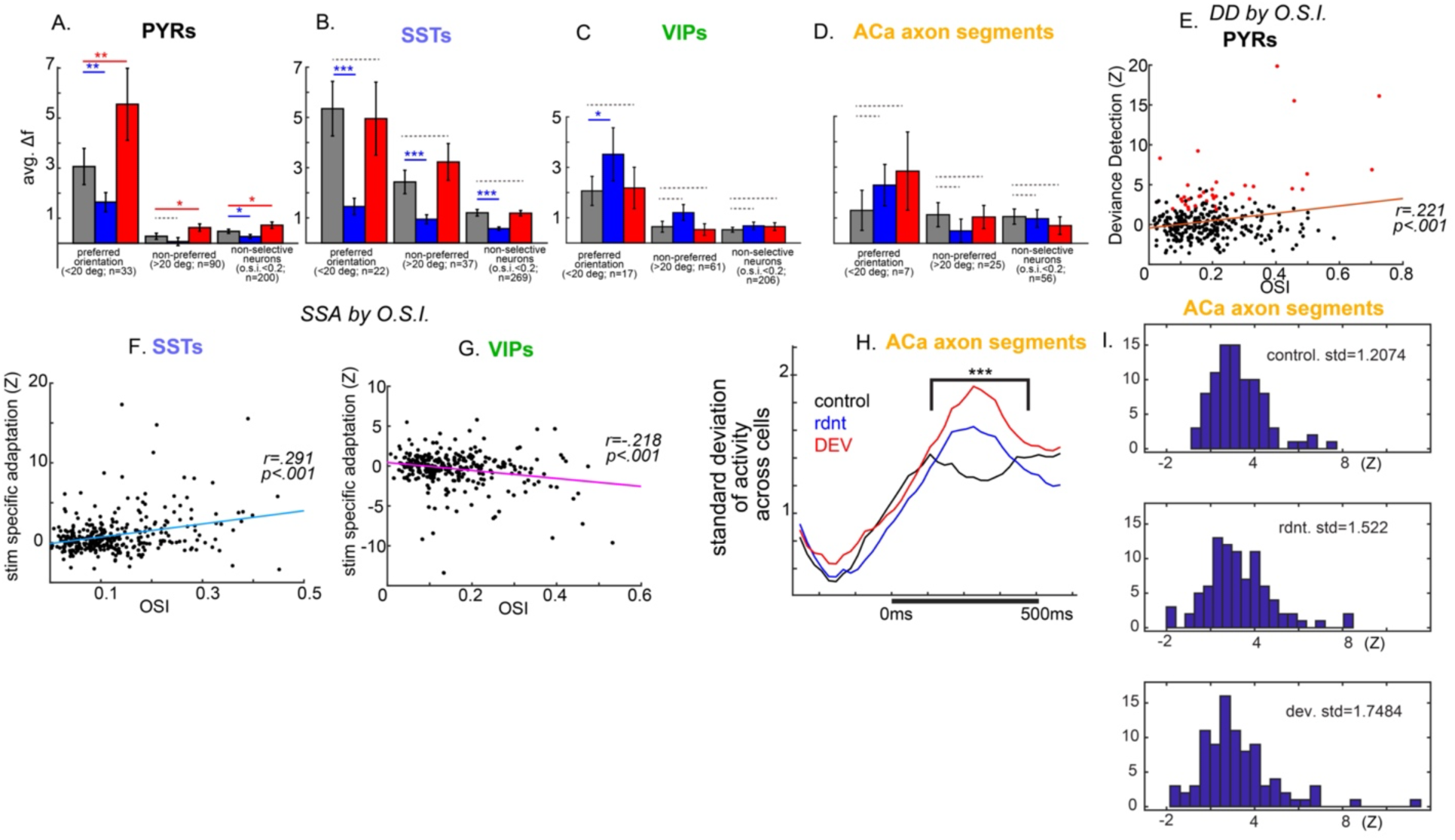
Deviance detection and stimulus specific adaptation by feature selectivity by cell-type. A) Pyramidal neurons displayed statistically significant deviance detection (DD) regardless of the cells preference for the orientation of the stimulus (one-tailed paired t-test, t-preferred (31)=2.63, p<.01; t-non-preferred(88)=1.82, p<.05), and in cells with low/absent orientation selectivity (t-non-selective (198)=1.81, p<.05). For stimulus specific adaptation (SSA), PYR responses to their preferred orientation (one-tailed paired t-test, t-preferred (31)=-2.93,p<.01) or non-selective PYRs (t-non-selective (198)=-1.86, p<.05) showed significant SSA. The lack of SSA to non-preferred stimulus in PYRs was likely due to floor effects, as responses to the control stimuli were already quite low. B) Neither selective nor non-selective SSTs showed significant DD, but all three groups showed significant SSA (one-tailed paired t-test, t-preferred (20)=-3.91, p<.001; t-non-preferred(35)=-3.72, p<.001; t-non-selective (297)=-5.21, p<.001). C) Neither selective nor non-selective VIPs showed significant DD or significant SSA. Interestingly, up examination, selective VIPs showed statistically significant inverse SSA to their preferred orientation (two-tailed paired t-test, t-preferred (15)=2.19, p<.05) and trend-level inverse SSA to their non-preferred orientation (t(61)=1.62, p=.11). D) Axonal segments from ACa neurons projecting to V1, did not show significant DD or SSA, regardless of the cells selectivity or stimulus preference. E) DD in PYRs was modestly but significantly correlated with orientation selectivity (1-circ variance). Still, many cells showing low orientation selectivity showed reliable DD (red cells in plot). DD was not correlated with orientation selectivity in SSAs, VIPs, or ACa axonal segments. F) SSA in SSTs was modestly but significantly correlated with orientation selectivity (a similar effect was observed in PYRs, not shown), while G) SSA in VIPs was inversely correlated with orientation selectivity, suggesting that more selective VIP neurons actually produced stronger responses to predictable stimuli. H,I) Although ACA axons in V1 do not show differences in average activity levels between control and oddball paradigms (see Fig 3 in text), they do exhibit increased standard deviation across population responses during the oddball paradigm, during both redundant and deviant trials, relative to control. This suggests that the overall nature of information being sent to V1 during the oddball paradigm is different than during the control.

**Figure S3.**
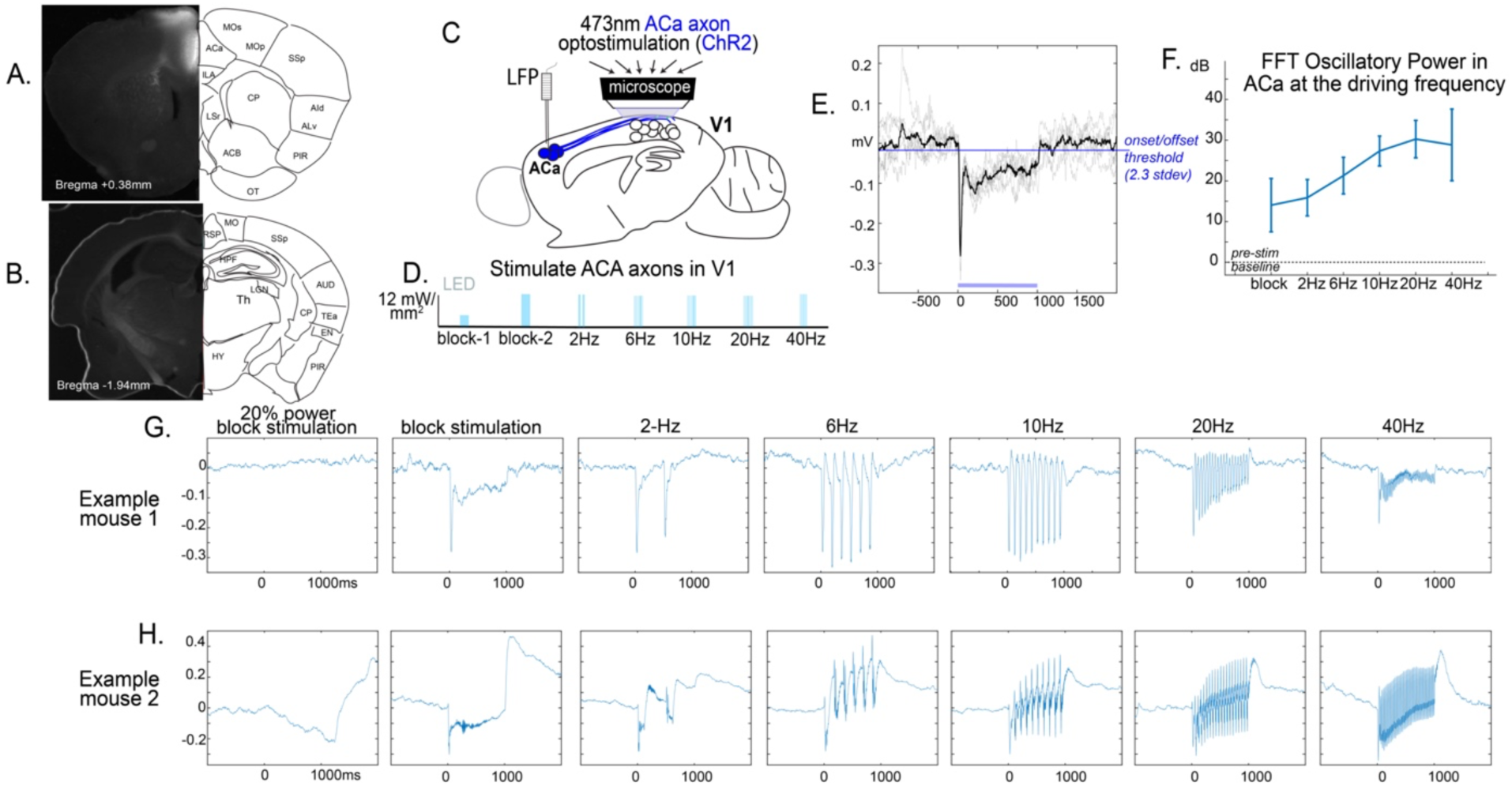
Electrophysiological confirmation of optogenetic driving. A) mCherry signal indicating expression of ChR2 in ACa neurons and B) not in areas posterior to ACa which project to V1. C) Axon terminals of long-range projecting neurons from ACa were stimulated in V1 via optogentics. Channel rhodopsin (ChR2) was expressed virally in ACa via AAV-9s through the synapsin promoter. L.E.D. powered 473 nm light focused (.5mm radius; 12 mW per mm^2^) via a 20x objective on a craniotomy was driven at D) 2-40 Hz (20% duty cycle) or block stimulation. Overall average power per 1 second was held constant for all conditions except the full power block stimulation. E) Local field potentials (LFP) were recorded in ACa to detect antidromic driving. On average (thick line), our stimulation evoked LFP potentials crossing 2.31 standard deviations above baseline (p<.01) at approximately 11.5 ms after light onset and returned to baseline at approximately 24.5 ms. Given known onset kinetics of ≈1.5 ms and offset kinetics of ≈ 13.5 ms (Lin, J.Y., Exp. Physiol 2012), we estimated a conduction speed of 10 ms from ACa to V1. F) Stimulus induced average power and standard deviations (across trials) display strong activation of ACa at the driving frequencies via illumination of the terminals. Notably, frequencies 6-40Hz are within 1 stdev of each other (n=2 mice, 8-15 trials each frequency per mice). G-H) Shows an averaged evoked LFP responses to each frequency for two representative mice. Notably, the low-power block stimulation was insufficient to drive ACa neurons in both mice.

**Figure S4.**
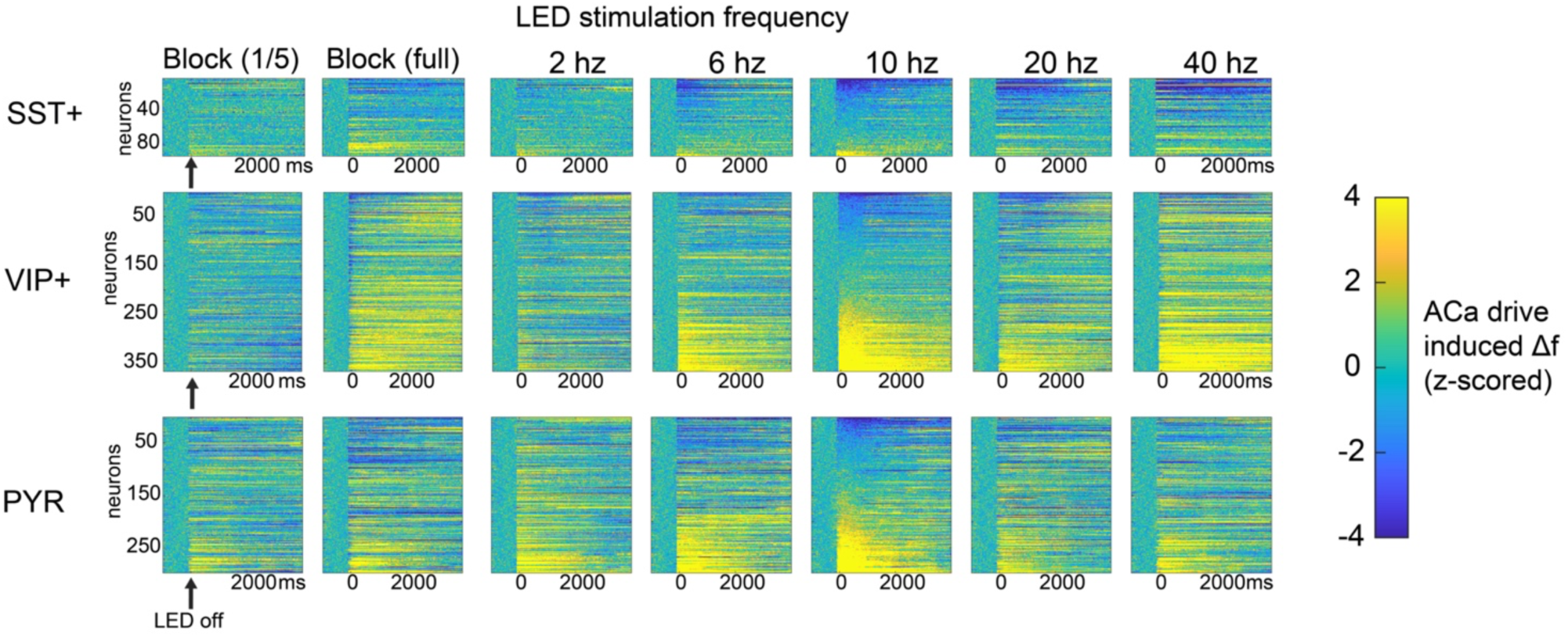
Activation of individual SST, VIP, and PYR neurons while driving ACa-axons at different frequencies. Rasterplots of activity immediately before turning on the LED and immediately after turning off the LED (0ms onward). Image signal was saturated during the 1 second of stimulation, so this data was excluded, and the 1 second post stimulation was used for analyses (figure 4). Activity of individual neurons was standardized by the standard deviation of the fluorescent signal in the 1-second of data prior to LED in order to allow for visual comparison across cells. Cells are sorted by their response to the 10-Hz condition. Cell identities are the same across all 7 conditions horizontally. Based on the large amount of variability across cells within each class, we carried out a cluster analysis to sort into functionally defined subclasses of neurons. See figure 4.

**Figure S5.**
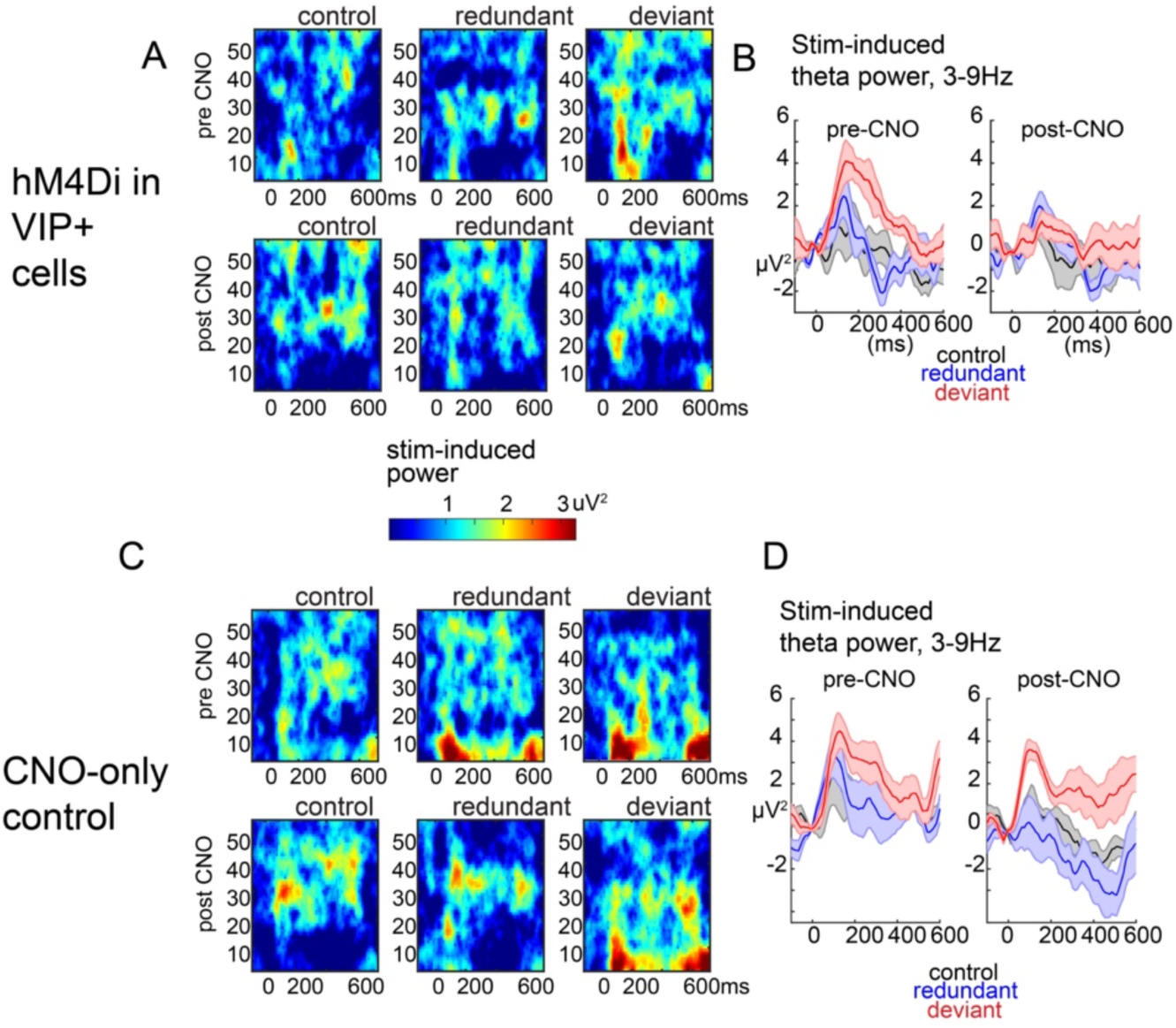
Chemicogenetic suppression of VIPs. A) time-frequency and B) line plots of induced power for pre- (above) and post-CNO treatment in mice with inhibitory DREADDs (hM4D(i)) in VIP interneurons. C,D) same as A,B, but for the CNO-only condition (no-DREADDs in VIPs).

**Figure S6.**
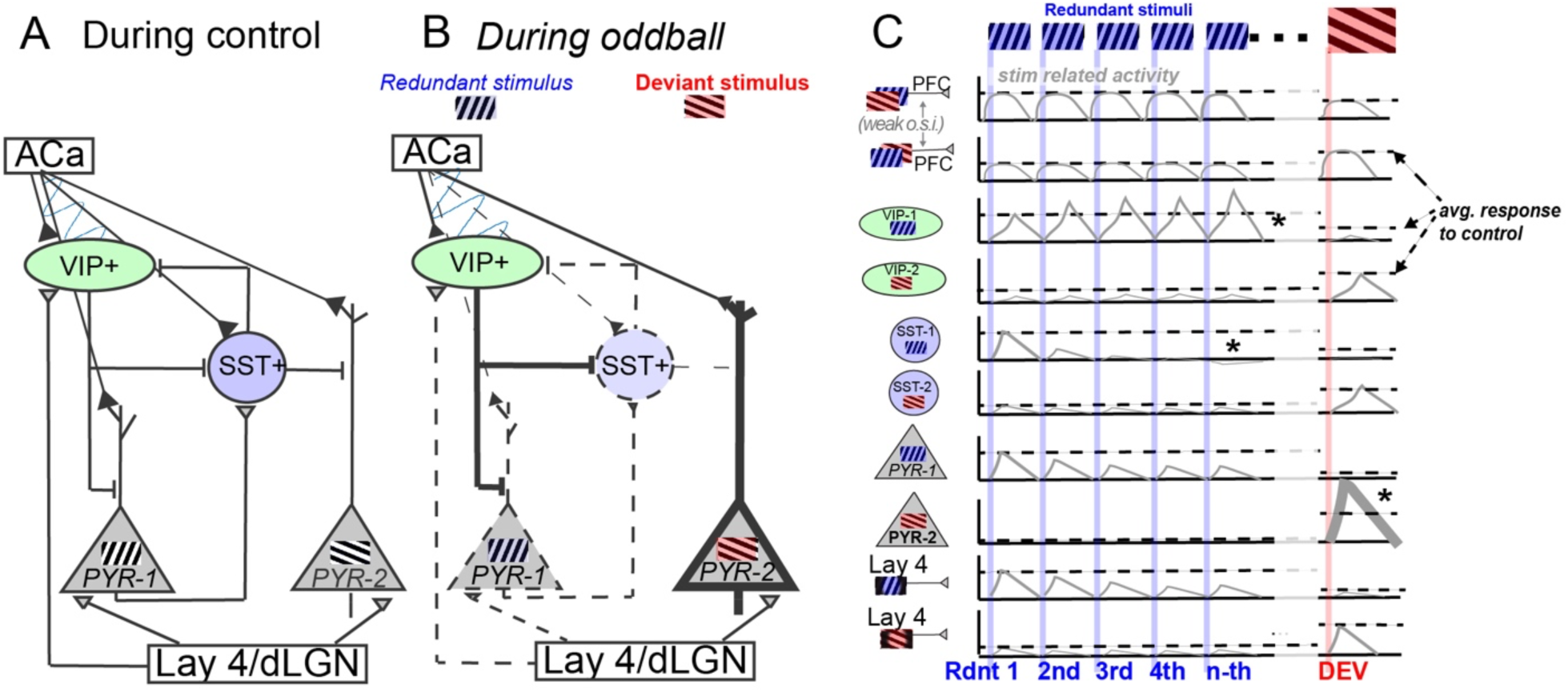
Proposed model of deviance detection in V1. A schematic based on current and past observations (with some testable assumptions about connectivity) to facilitate future work into deviance detection circuits. (A) simplified V1 layer 2/3 circuitry. During control run, with stimuli of many orientations equally likely, responses to any given stimulus by the circuit is at a baseline, with ACa and V1 synchronous in the theta/alpha band. B) During the oddball paradigm, inputs from ACa to V1 become more influential (as evidenced by greater theta/alpha granger causality) and more varied, potentially dependent on postsynaptic target (VIPs vs SSTs vs PYRs selective for redundant or deviant). Line-thicknesses depict how a cell population’s excitability is modulated during the oddball, relative to the many-standards control (thicker = greater response to its preferred stimulus over baseline). Top-down modulation at theta/alpha frequencies potentiates VIPs and suppresses SSTs, which, in turn, disinhibits PYRs which are not already adapted (i.e. PYRs selective for stimuli other than the redundant). (C) Activity dynamics during the oddball paradigm for each cell type, relative to its selectivity (based on figure S2). Horizontal dotted lines depict response of cell population to each stimulus orientation in the control context. From top: *absent SSA in ACA, *enhanced SSA in SST-1s to the redundant stimulus, and *DD in SST-2 and PYR-2. Layer 4 cell activity based on Hamm et al 2021^42^

